# Characterization of hairpin loops and cruciforms across 118,065 genomes spanning the tree of life

**DOI:** 10.1101/2024.09.29.615628

**Authors:** Nikol Chantzi, Camille Moeckel, Candace S. Y Chan, Akshatha Nayak, Guliang Wang, Ioannis Mouratidis, Dionysios Chartoumpekis, Karen M. Vasquez, Ilias Georgakopoulos-Soares

## Abstract

Inverted repeats (IRs) can form alternative DNA secondary structures called hairpins and cruciforms, which have a multitude of functional roles and have been associated with genomic instability. However, their prevalence across diverse organismal genomes remains only partially understood. Here, we examine the prevalence of IRs across 118,065 complete organismal genomes. Our comprehensive analysis across taxonomic subdivisions reveals significant differences in the distribution, frequency, and biophysical properties of perfect IRs among these genomes. We identify a total of 29,589,132 perfect IRs and show a highly variable density across different organisms, with strikingly distinct patterns observed in Viruses, Bacteria, Archaea, and Eukaryota. We report IRs with perfect arms of extreme lengths, which can extend to hundreds of thousands of base pairs. Our findings demonstrate a strong correlation between IR density and genome size, revealing that Viruses and Bacteria possess the highest density, whereas Eukaryota and Archaea exhibit the lowest relative to their genome size. Additionally, the study reveals the enrichment of IRs at transcription start and termination end sites in prokaryotes and Viruses and underscores their potential roles in gene regulation and genome organization. Through a comprehensive overview of the distribution and characteristics of IRs in a wide array of organisms, this largest-scale analysis to date sheds light on the functional significance of inverted repeats, their contribution to genomic instability, and their evolutionary impact across the tree of life.

## Introduction

DNA is the genetic information carrier molecule used by most organisms. Since the discovery of the DNA double-helix structure (B-DNA structure) over 70 years ago, where two antiparallel strands intertwine, many different types of alternative conformations (i.e. non-B DNA) that differ from the classic B-DNA conformation have been discovered. These non-B DNA structures play a wide variety of biological roles in genomic DNA organization, replication, transcription, and recombination and may provide positive selection advantages that make certain non-B DNA motifs conserved (Kaushik et al. 2016; Makova and Weissensteiner 2023). On the other hand, many types of non-B DNA structures are intrinsically mutagenic and therefore provide driving forces for genome evolution and disease development (Wang and Vasquez 2022; Georgakopoulos-Soares et al. 2018; Cheloshkina and Poptsova 2019).

Alternative DNA conformations can be inferred from the primary nucleotide sequence. One of these conformations is the hairpin/cruciform DNA structure, which forms at inverted repeat (IR) sequences or palindromic motifs (Bikard et al. 2010; Buisson et al. 2019; Lu et al. 2015; Bacolla et al. 2016) (**Figure 1A**). In these instances, double-stranded DNA adopts hairpin conformations, with a single DNA strand with two symmetric sequences undergoing intrastrand base pairing to form hairpin or cruciform structures consisting of two complementary arms separated by a spacer loop (**Figure 1A**). Physiological processes such as DNA transcription and replication, which involve the opening of double-stranded DNA into a single-stranded state, generate negative supercoiling to facilitate hairpin and cruciform formation (Azeroglu et al. 2014; Rosche, Trinh, and Sinden 1995).

**Figure 1:**
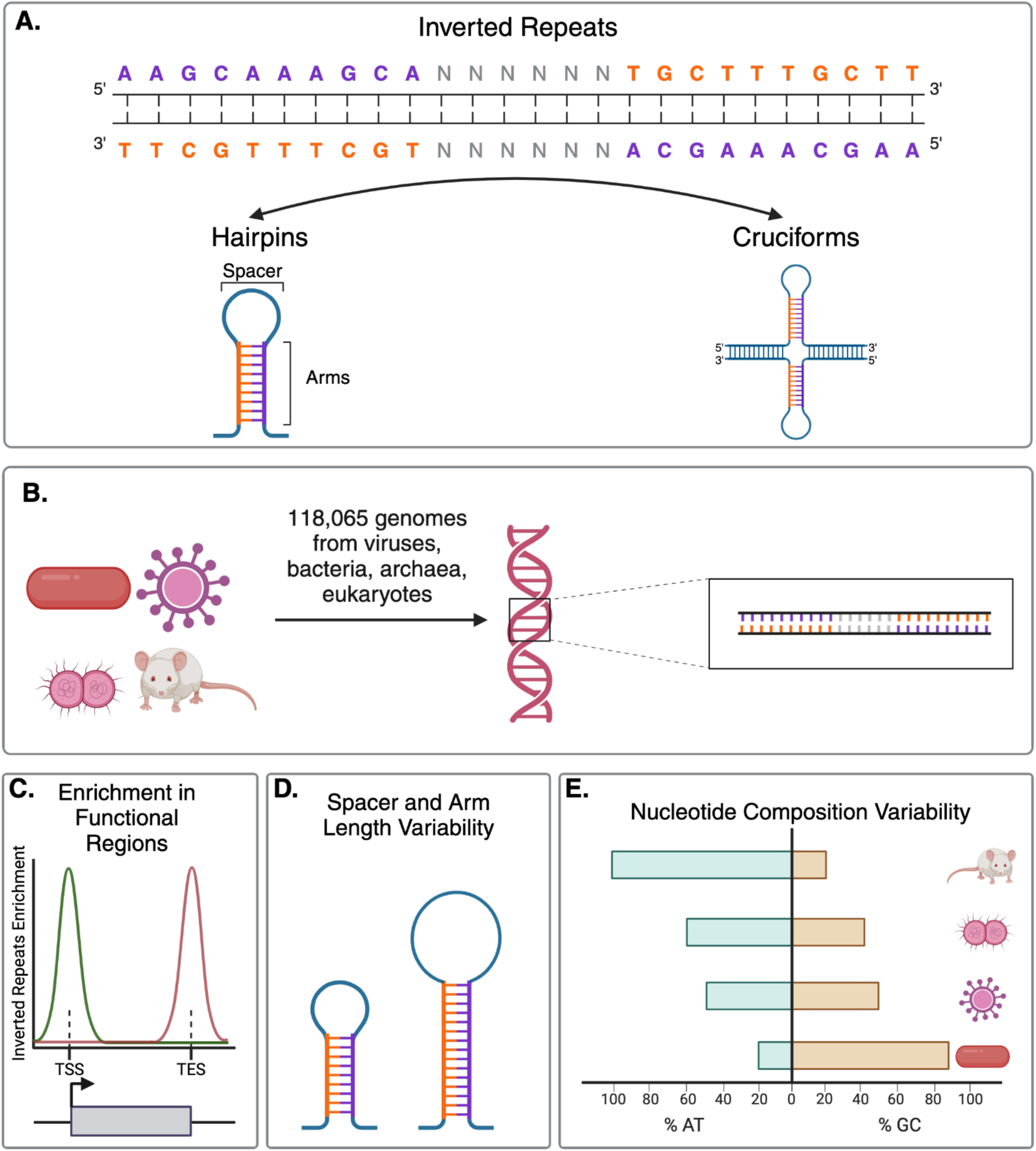
Schematic representation of the investigation of IRs across the tree of life in this study. **A.** Schematic representation of an inverted repeat. Colored base-pairs denote the IR arms, and the “N” letters represent the intervening spacer region. Formation of hairpin and cruciform structures can occur at IRs. **B.** IR identification was performed across 118,065 complete organismal genomes, including Viruses, Bacteria, Archaea, and Eukaryota. **C.** We observed enrichment of IRs relative to functional genomic loci, including transcription start sites and transcription end sites. **D-E.** We found that biophysical properties and the nucleotide composition of IRs, including spacer and arm length, influence their frequency across taxa.

The biophysical properties of IRs impact the likelihood of hairpin/cruciform structure formation. Stable hairpin/cruciform formation is promoted by longer IR arms, while mismatches in the hairpin arms and longer spacer lengths reduce the likelihood of stable structure formation (Dionisios Rentzeperis et al. 2002; Nag and Petes 1991; Nasar, Jankowski, and Nag 2000; K. S. Lobachev et al. 1998; Sinden et al. 1991). Additionally, the GC content of the stem is associated with increased thermodynamic stability of the DNA hairpin/cruciform (Buisson et al. 2019; Woodside et al. 2006), and the cellular environment contributes to the likelihood of hairpin formation (Tan and Chen 2008).

IRs contribute to genetic instability across prokaryotic and eukaryotic organisms (Gordenin et al. 1993; K. S. Lobachev, Rattray, and Narayanan 2007; Nag and Kurst 1997; K. S. Lobachev et al. 1998; Butler, Gillespie, and Steele 2002; Achaz et al. 2003; Leach 1994; Tanaka et al. 2002; Zhou, Akgūn, and Jasin 2001; Lindsey and Leach 1989). In particular, maintaining long IRs *in vivo* poses a significant challenge (Leach 1994; K. S. Lobachev et al. 1998; Nag and Kurst 1997). In the human genome, IRs serve as mutational hotspots, accumulating an excess of both germline and somatic mutations (Langenbucher et al. 2021; Buisson et al. 2019; Georgakopoulos-Soares et al. 2018; Guiblet et al. 2021; Lu et al. 2015; Zou et al. 2017; Georgakopoulos-Soares, Victorino, et al. 2022; Bacolla et al. 2016). As a result, the vast majority of IRs are less than 100 base-pairs (bps) long (H. Kurahashi and Emanuel 2001; Wang and Vasquez 2006). Longer IR sequences do exist in the human genome, particularly in the X and Y chromosomes, but they tend to contain multiple mismatches (Warburton et al. 2004). Long (>500bp), AT-rich IRs with mismatches have been associated with gross chromosomal rearrangements in the human genome (H. Kurahashi and Emanuel 2001).

The rapid growth of available genomic data in recent years provides a unique opportunity to study genome structure and composition comprehensively across a wide range of organisms from different taxa of the tree of life. In the coming years, the number of complete organismal genomes is expected to increase exponentially and encompass a significant proportion of the genetic diversity present in nature. For example, the Earth BioGenome Project aims to sequence the genomes of all eukaryotic species within the next ten years (Lewin et al. 2022). Analyzing the diversity of organismal genomes is essential for uncovering biological insights and has far-reaching implications for advancing our understanding of evolutionary history, unraveling the complexities of functional genomics, and improving our understanding of human health and disease. Previous research on 1,565 bacterial genomes revealed that, for IRs with arm lengths of six bps or longer, the highest and lowest mean frequencies were observed in the phylum Tenericutes (also known as Mycoplasmatota) and the class Alphaproteobacteria, respectively (Porubiaková et al. 2023). Another study examining promoters from 1,180 organismal genomes found an enrichment of IRs in the promoters of prokaryotic organisms (Yella and Vanaja 2023). IRs have also been systematically characterized in chloroplast and mitochondrial genomes (Cechová et al. 2018; Brázda et al. 2018).

Functional roles for IRs have been uncovered (Bikard et al. 2010; Brázda et al. 2011; Grechishnikova and Poptsova 2016). For example, IRs are found at replication origins in prokaryotes, eukaryotes, and viruses (Pearson et al. 1996; Leung et al. 2005) and play diverse roles in gene expression regulation. In gene promoters, they act as inhibitory elements and modulate transcription termination, but at the RNA level, they influence steady-state mRNA expression and stability in 3′ untranslated regions (Georgakopoulos-Soares, Victorino, et al. 2022; Siegel et al. 2022; Brázda et al. 2020; de Hoon et al. 2005; Chen et al. 2013; Fleming et al. 2020; Georgakopoulos-Soares, Chan, et al. 2022; Miura, Ogake, and Ohyama 2018). In prokaryotes, IRs are pivotal structures in rho-independent transcription termination (Kingsford, Ayanbule, and Salzberg 2007; von Hippel 1998). Additionally, research has identified associations between specific proteins and hairpin or cruciform structures (Brázda et al. 2011). Despite these findings, no comprehensive study to date has systematically characterized IRs across organisms throughout the entire tree of life.

Here, we perform a comprehensive investigation of perfect IRs (without mismatches in the arms, with minimum arm lengths of ten bps, and with spacer lengths up to eight bps) across 118,065 complete organismal genomes for the domains of life and for Viruses. We report a total of 29,589,132 IRs and find that the density of IRs per genome varies significantly among different phyla. Viruses and Bacteria harbor the highest density of IRs relative to their genome size, whereas Eukaryota and Archaea have the lowest. Genome size is positively correlated with IR density in Archaea and Eukaryota but negatively correlated in Bacteria and Viruses. Nevertheless, the density of IRs per genome varies significantly among different phyla. We also find that the biophysical properties of IRs are highly biased between taxonomies. Eukaryotic IR arms are highly enriched for repetitive AT-rich sequences. Notably, we observe that *Plasmodium falciparum* has the highest IR density among organisms from the domains of life studied, and these IRs are located in highly AT-rich, tandemly repeated regions. Bacteria show a preference for spacers of four bps, whereas Eukaryota give rise to no-spacer perfect palindromes due to the inherently repetitive organization of eukaryotic genomes. Finally, we find that IRs are preferentially positioned upstream of Transcription Start Sites (TSSs) and downstream of Transcription End Sites (TESs), and Bacteria have a strong preference (exceeding 12-fold enrichment) for IRs downstream of TESs. We conclude that perfect IRs display large discrepancies in abundance between organismal genomes, have notable biophysical differences between taxonomies, and are preferentially located in functional elements.

## Results

### Identification of perfect IRs across 118,065 organismal genomes

To determine the frequency of long, perfect IRs in nature across organismal genomes, we used 118,065 complete genomes from organisms spanning taxonomic subdivisions. We focused on perfect IRs, those without mismatches in the arms, as they are more likely to form hairpin/cruciform structures. We used arm sizes longer than nine bps and spacer lengths up to eight bps. These limits were used because IRs with short arms and longer spacer loops are less likely to fold into hairpin/cruciform structures (Dionisios Rentzeperis et al. 2002; Nag and Petes 1991; Nasar, Jankowski, and Nag 2000; Sinden et al. 1991).

We mapped out IR sequences across the genomes of each organism for which sequencing data was available, creating a comprehensive genome-wide dataset with species from diverse taxa across the tree of life. We found 29,589,132 IRs. Upon examining the IR densities across the distinct taxonomic partitions, we note that Bacteria exhibit the highest average IR density of 3,253.64 IR bps per megabase (Mb), followed by Viruses and Eukaryota (**Figure 2A-C**). However, out of 67,725 examined viruses, 33,142 exhibited no IRs with the aforementioned parameters. We observed remarkable variation in the density of IRs across organismal genomes, with densities ranging from 0 IR bps per Mb to 180,327.86 IR bps per Mb, and a median density of 1,782.44 IR bps per Mb. The maximum IR density was attained for *Kummerowia striata partitivirus* (**Figure 2A**; **Supplementary Figure 1**).

**Figure 2:**
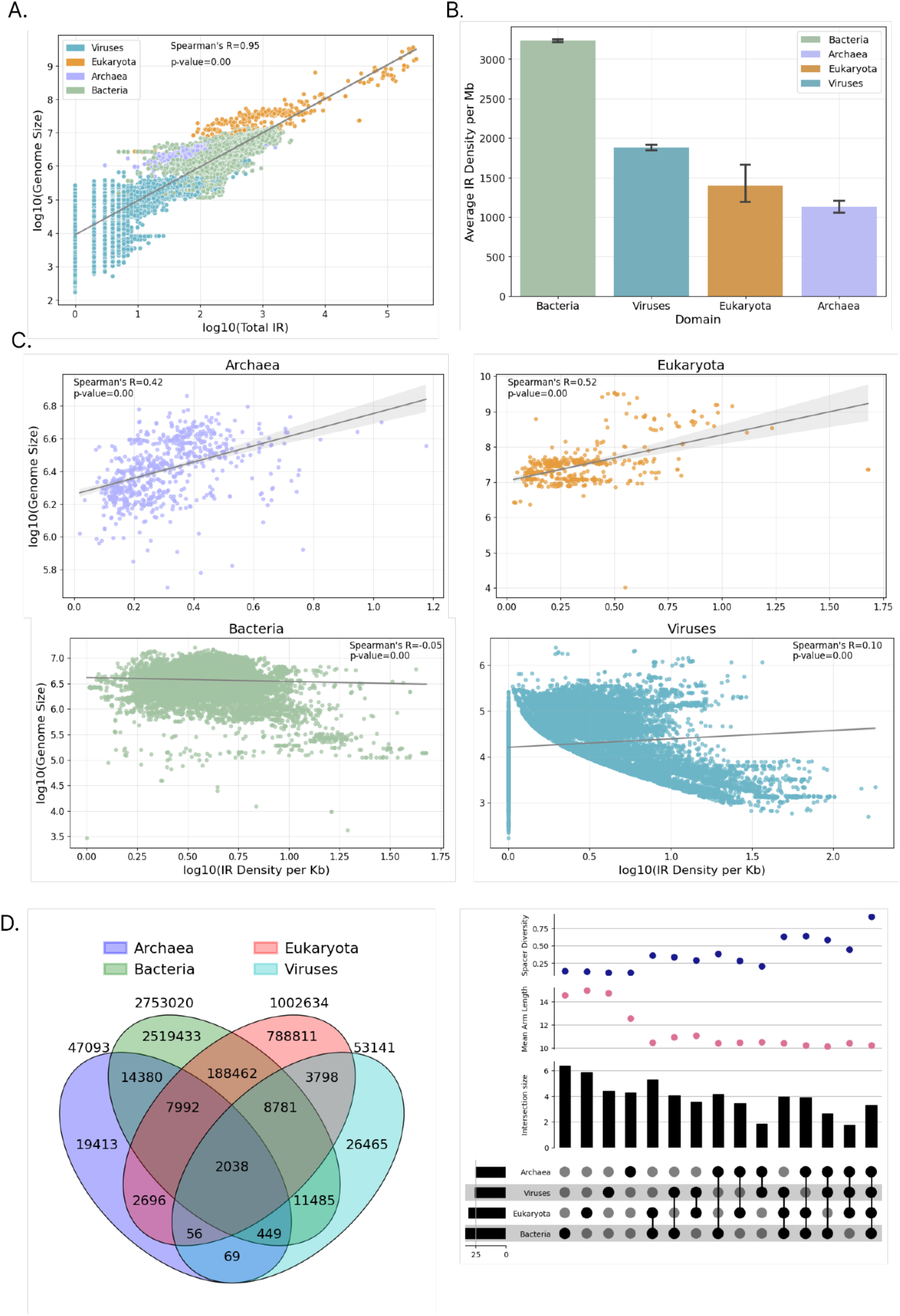
Characterization of IRs in 118,065 organismal genomes across the three domains of life and Viruses. **A.** Association between genome size and the number of IRs identified. Each dot represents an organismal genome, with dot color indicating one of the three domains of life or Viruses. **B.** Average density of IR bp per Mb for organisms in the three domains of life and Viruses. Error bars represent standard error. **C.** Association between genome size and the proportion of the genome covered by IRs, presented separately for the three domains of life and Viruses. The correlation for Viruses excludes genomes without IRs. **D.** Representation of the number of shared sequence arms for IRs identified in organismal genomes across the three domains of life and Viruses. This includes a Venn diagram and an upset plot representing the shared number of IR arms and the proportions shared between the three domains of life and Viruses. Spacer density and arm density are also displayed for each comparison.

We were interested in finding the longest IRs present in organismal genomes; therefore, we investigated whether there are IRs of hundreds or thousands of bps in any of the studied genomes. Although rare, we found several IRs with arm lengths of hundreds or tens of thousands of bps (**Table 1**; **Supplementary Figure 2**). In particular, the largest IR, with an arm length of 117,051 bp, was detected in the bacterium *Klebsiella pneumoniae* on chromosome NZ_CP116903.1, encompassing a large area of 226 protein-coding genes. However, IRs with perfect arms of extreme lengths were not identified in *Homo sapiens*, with the longest IR detected having arms of 140 bp (**Supplementary Table 1**). We conclude that IRs of arm lengths of thousands of bps without mismatches are almost exclusively found in Bacteria.

**Table 1:** Longest, perfect IRs across the genomes studied.

| Inverted Repeat Arm Length (bps) | Genome | Spacer Length (bps) | Arm GC Content (%) |
| --- | --- | --- | --- |
| 117,051 | *Klebsiella pneumoniae* | 1 | 56.455 |
| 52,115 | *Enterococcus faecium* | 0 | 33.385 |
| 47,461 | *Enterococcus faecium* | 0 | 32.814 |
| 38,727 | *Olsenella timonensis* | 0 | 69.365 |
| 38,727 | *Atopobiaceae bacterium* | 0 | 69.365 |
| 38,611 | *Rhodococcus sp. AH-ZY2* | 1 | 64.945 |
| 38,528 | *Pseudomonas aeruginosa* | 0 | 66.824 |
| 37,277 | *Pseudomonas corrugata* | 0 | 60.627 |
| 30,325 | *Klebsiella pneumoniae* | 0 | 51.023 |
| 27,900 | *Enterococcus faecium* | 0 | 33.225 |
| 22,733 | *Staphylococcus pasteurii* | 1 | 30.625 |
| 18,812 | *Ligilactobacillus murinus* | 0 | 35.461 |
| 18,297 | *Escherichia coli* | 0 | 45.335 |
| 17,045 | *Acanthamoeba polyphaga mimivirus* | 0 | 26.617 |
| 14,452 | *Agrobacterium pusense* | 0 | 55.272 |
| 13,509 | *Melissococcus plutonius* | 0 | 29.350 |
| 13,040 | *Salmonella enterica* | 4 | 49.095 |

### Discrepancies in IR frequencies in organismal genomes across the tree of life

We investigated IRs across the three domains of life and Viruses separately. Interestingly, we observed significant differences in the frequency of perfect IRs between Archaea, Eukaryota, Bacteria, and Viruses. Bacteria and Viruses exhibited the highest genomic density of IRs with an average density of 3,231.28 and 1,877.80 IR bps per Mb, respectively, whereas Eukaryota and Archaea displayed the lowest IR densities per genome with 1,393.59 and 1,129.86 IR bps per Mb, respectively (**Figure 2D**). Notably, we found that the proportion of the genome covered by IRs is not strongly correlated with genome size in Viruses (**Figure 2C**, Spearman correlation r = 0.1, p-value=0). For Archaea (Spearman correlation r=0.42, p-value=0.0) and Eukaryota (Spearman correlation r=0.52, p-value=0.0), this proportion is positively correlated with genome size (**Figure 2C**). We also observed that Eukaryota display a high level of repetitiveness in IR arm sequences and are enriched for AT-rich sequences, which are also associated with tandem repeats. Finally, in Bacteria, we found a weak negative correlation between IR genome coverage and genome size (**Figure 2C**, Spearman correlation r=-0.11, p-value=7.2e-98). These findings suggest differences in the frequency of IRs associated with taxonomic groups and genome size.

To gain insights into differences in the composition of IRs across taxonomies, we examined the number and proportion of IR arm sequences appearing only in one of the three domains of life or Viruses and not in the others. Bacteria displayed a high proportion of arm sequences that were unique and not found in the other taxonomies compared to the other three domains (**Figure 2D**; **Supplementary Figure 3**). Moreover, a significant proportion of viral IR arm sequences are shared between the viral and bacterial domains (**Figure 2D**). We also observed that IR arm sequences shared between domains display a higher average diversity in spacer length, indicating that these omnipresent IR arm sequences appear with many different spacer length variations compared to sequences exclusive to one, two, or all three domains (**Figure 2D**). Thus, we conclude that the spacer length of IR arms unique to Bacteria is more likely to be short. Finally, the mean arm size peaks in the IRs originating solely from the eukaryotic organismal genomes.

### Substantial differences in the IR spacer and arm length distributions among taxa

The biophysical properties of IRs, including the spacer and arm lengths, as well as the nucleotide composition of the hairpin arms, influence the likelihood of hairpin/cruciform formation and its stability (Nag and Petes 1991; Varani 1995). Therefore, we examined the effect of these biophysical properties on the frequencies of IRs across taxonomic subdivisions. We observed that across taxonomic groups the largest number of IRs have short arm lengths (**Figure 3A**), which is expected since the probability of finding short complementary sequences by random chance is higher than for longer sequences. In contrast, we find that the number of IRs identified varies significantly for different spacer lengths between different taxonomic groups. In Eukaryota, IRs are predominantly found with zero spacer lengths, whereas in Bacteria, the largest number of IRs have spacer lengths of four bps (**Figure 3B-C**). Archaea show smaller differences between spacer lengths but tend to prefer shorter spacers, whereas in Viruses, there is a preference for spacer lengths of zero bps (**Figure 3B-C**).

**Figure 3:**
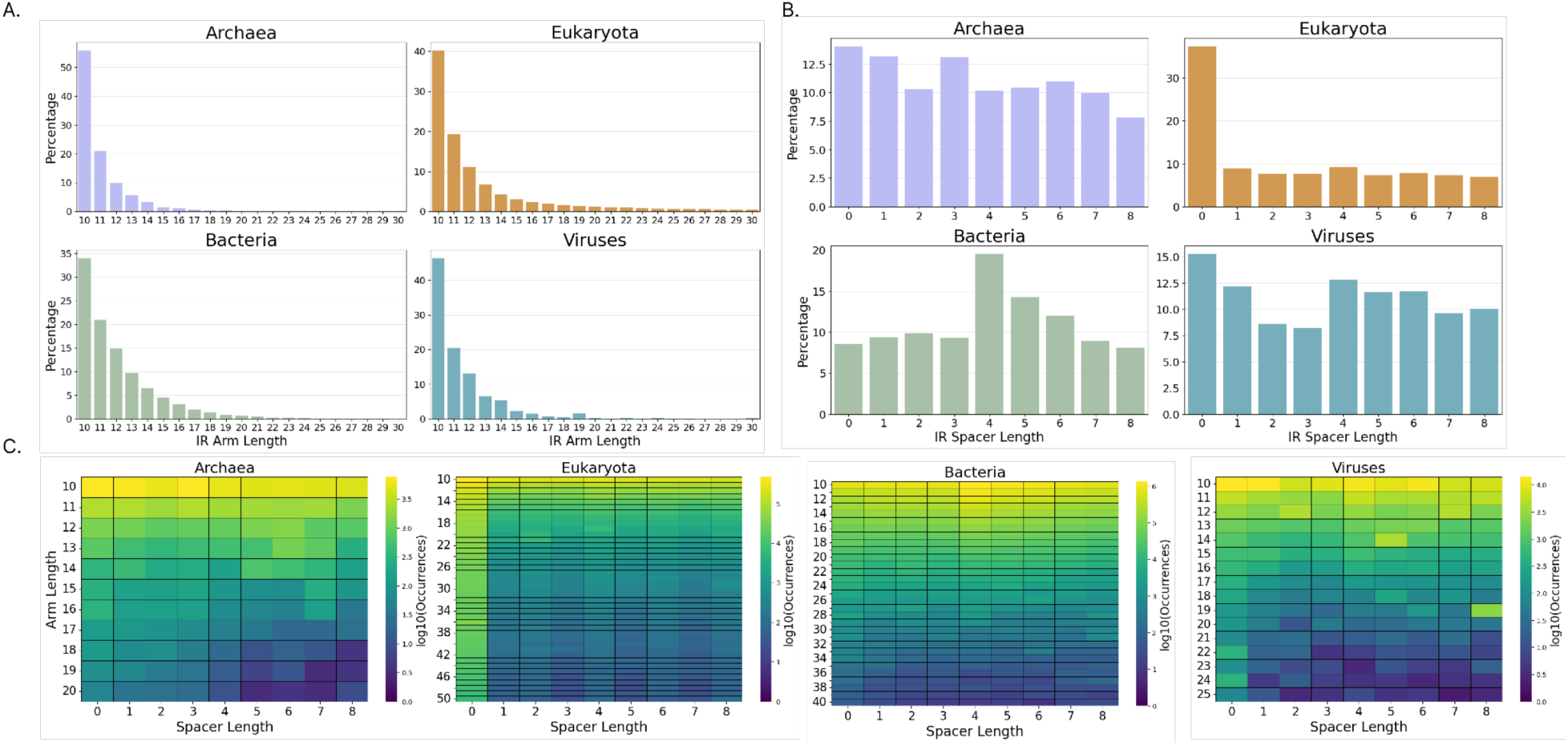
Distribution and density of IR types across taxonomic subdivisions. **A.** Number of IRs as a function of arm length, and **B.** Percentage (%) of IRs with corresponding spacer length, across the three domains of life and Viruses. **C.** Heatmaps showing the occurrences of IRs as a function of arm and spacer length combinations.

Next, we investigated differences in the genomic density of IRs across kingdoms and phyla. The kingdoms with the highest coverage were eukaryotic and included Protista, Plantae, and Animalia, with 3,992.32, 3,476.28 and 3,062.79 IR bps per Mb, respectively. These IRs tended to be AT-rich and often overlapped tandem repeats. Eubacteria followed closely with 2,891.40 IR bps per Mb, while viral kingdoms, such as Shotokuvirae with 2,981.09 IR bps per Mb, Sangervirae with 2,962.34 IR bps per Mb, and Loebvirae with 2,634.75 IR bps per Mb also displayed notable densities (**Figure 4A**). In contrast, Fungi exhibited the lowest IR density with 857.81 IR bps per Mb. These findings indicate significant differences in the IR genomic content across kingdoms, even within the same domain. At the phylum level, the two phyla with the highest density of IRs were both bacterial, namely *Candidatus Campbellbacteria* and *Candidatus Gracilibacteria* with densities of 11,546.31 and 9,561.79 IR bps per Mb, respectively (**Figure 4B**; **Supplementary Figure 4**). Additionally, the *Apicomplexa*, a protist phylum, had an IR genomic density of 6,350.21 bps per Mb. High IR densities were also observed in *Candidatus Kaiserbateria*, *Mycoplasmatota*, *Streptophyta*, and *Chordata*, which belonged to Bacteria, Plants, and Animals (**Figure 4B**). These findings underscore the substantial variability in the IR density across domain, kingdom, and phylum levels.

**Figure 4:**
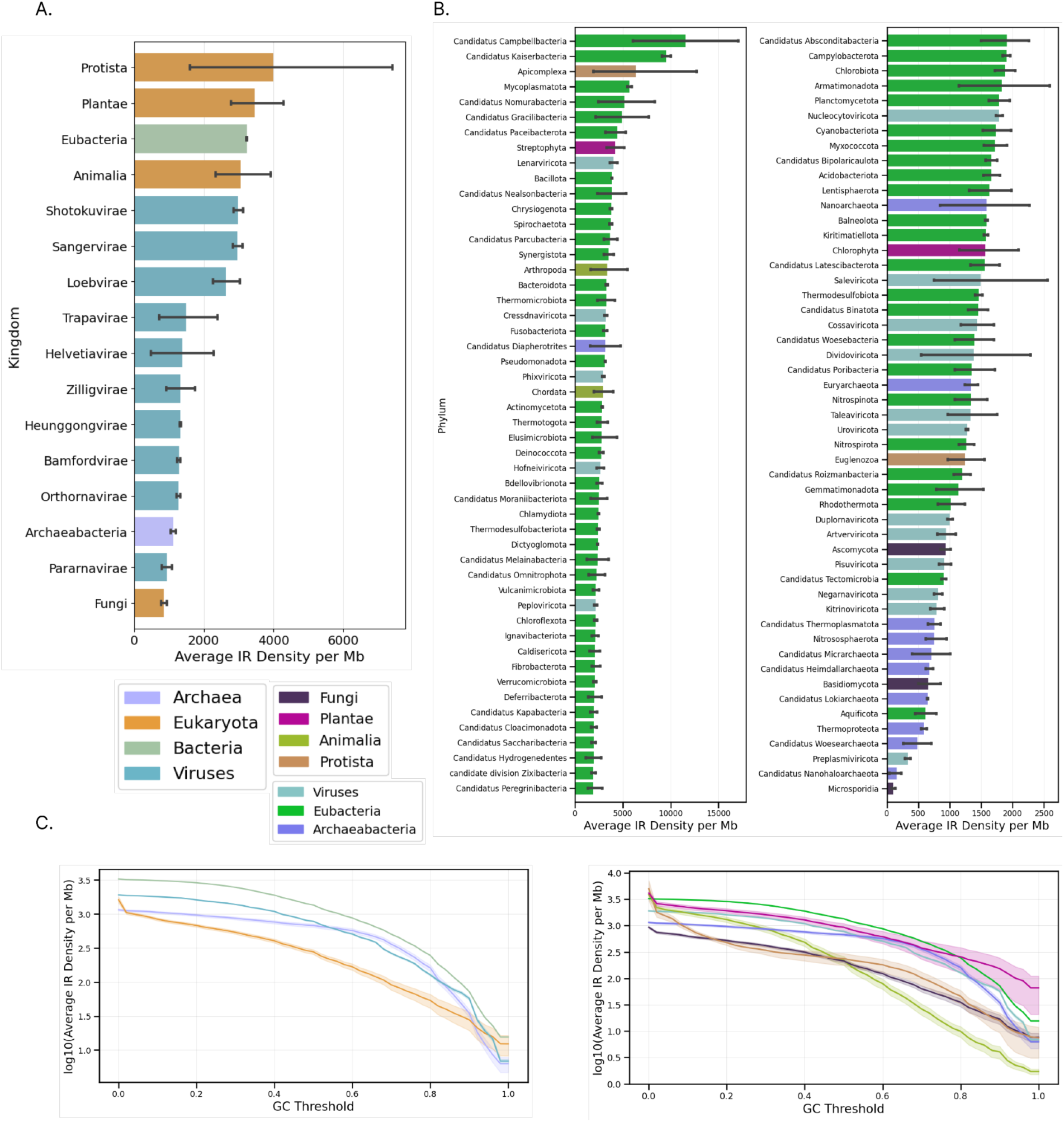
Genomic density of IRs in organismal genomes across the tree of life. **A.** Average IR coverage across organismal genomes belonging to the same kingdom. The color represents the three domains and Viruses to which each of the kingdoms belongs. **B.** Average IR coverage across different phyla. The color represents the kingdom to which each of the phyla belongs. Error bars represent standard error. **C.** IR Density Depletion relative to the GC Threshold of the arm sequence. The GC threshold is defined as the proportion of the GC content in the IR. Confidence intervals are calculated using the standard error of IR density in each organismal genome.

Next, we examined the contribution of AT content to the number of IRs detected per organism across kingdoms and phyla. We investigated the genomic density of IRs as a function of AT content. We observed that in Eukaryota, there is a steep decline in the number of IRs detected as we increase the percent of GC content in the arm (**Figure 4C**). On the other hand, such a sharp decline is not observed in Viruses, Bacteria, or Archaea, indicating that the vast majority of eukaryotic IRs are GC-depleted. This shows that when excluding AT-rich IRs, Eukaryota have the lowest IR density compared to Archaea, Bacteria, and Viruses and this pattern is consistent at the kingdom level (**Figure 4C**). Specifically, when examining IRs with a GC content of at least 10%, we find that the highest genomic density is observed in *Eubacteria*, followed by three viral kingdoms, namely *Sangervirae*, *Shotokuvirae*, and *Loebvirae*. This consistency is also observed for a GC content threshold of 20% (**Supplementary Figure 5**). This result is only partially accounted for by differences in the GC content of organismal genomes, indicating a subset of highly prevalent AT-rich and repetitive IRs in Eukaryota. We conclude that repetitive, AT-rich IRs are the most common IRs in Eukaryota, accounting for the vast majority of IR occurrences in Eukaryota, but not in Bacteria, Archaea, or Viruses.

### Biophysical properties of IRs across taxonomic groups in the tree of life

Next, we investigated the nucleotide composition of the arms relative to that of each organism’s genome across the three domains of life and Viruses. We observed an overall higher AT content in the IR arms of Eukaryota and Archaea relative to the overall AT content of their genome (**Figure 5B**). Additionally, we observed that in Eukaryota, there is higher AT content in the IR arms.

**Figure 5:**
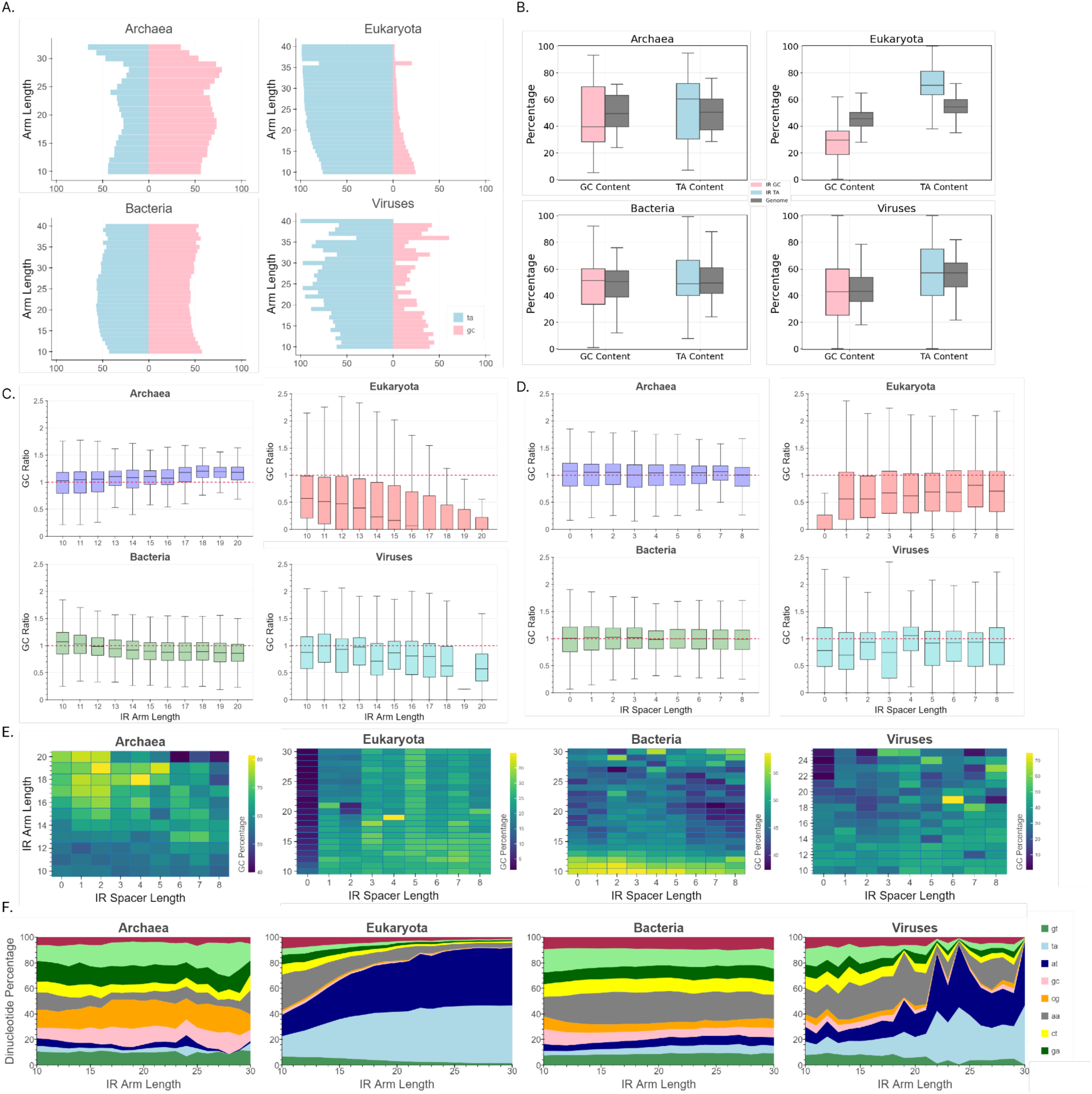
Characterization of IRs by their biophysical properties and nucleotide composition. **A.** Comparison of the GC and AT content in IR arms across genomes as a function of arm length. **B.** GC ratio in arms versus the genome.. **C-D.** GC ratio in arms versus the genome as a function of **C**. arm and D. spacer length. **E.** Heatmaps showing the GC content of IRs as a function of arm and spacer length combinations. **F.** Percentage of the different dinucleotides in arms as a function of arm length.

Viruses also displayed high AT content, which increased with IR arm length (**Figure 5A**). In contrast, such differences were not observed in Bacteria and Archaea, with Archaea showing an excess of GC relative to AT in the IR arms (**Figure 5A**).

When examining the GC content of the arms relative to that of each organism’s genome, we found a positive correlation in Archaea (Spearman ρ=0.84, p-value=0.0052 with Bonferroni corrected p-values), but a negative correlation in Bacteria (Spearman ρ=-0.97, p-value= 2.06e-06 with Bonferroni corrected p-values), Eukaryota (Spearman ρ=-0.92, p-value=7.089e-05, with Bonferroni corrected p-values), and Viruses (Spearman ρ=-0.78, p-value=0.0179, with Bonferroni corrected p-values) (**Figure 5B-D**). This suggests that shorter arms tend to have higher GC content in Eukaryota, Bacteria, and Viruses, but not in Archaea. When performing the same analysis separated by spacer length, we found that the main deviation is in Viruses and Eukaryota, in which IRs without spacers tend to be highly AT-rich compared to the genomic background (**Figure 5C-D**).

Next, we investigated the dinucleotide content of the arms as a function of IR length. We found that IRs with longer arm lengths have a disproportionate AT/TA content in Eukaryota (Spearman ρ=0.795, p-value=6.68e-4, with Bonferroni corrected p-values), Bacteria (Spearman ρ=0.854, p-value=3.26e-6, with Bonferroni corrected p-values), and Viruses (Spearman ρ=0.863, 1.83e-6, with Bonferroni corrected p-values) (**Figure 5E**), but not in Archaea (p-value>0.05). Therefore, we conclude that the biophysical properties of IRs, which influence the likelihood and stability of hairpin/cruciform formation, differ substantially between the three domains of life and Viruses.

### IRs are inhomogeneously distributed at functional genomic elements and are positioned relative to transcription start sites (TSSs) and termination end sites (TESs)

Previous work has shown that IRs are inhomogeneously distributed in the human genome and have a number of functional roles in a number of other organismal genomes (Georgakopoulos-Soares, Victorino, et al. 2022; Siegel et al. 2022; Brázda et al. 2020; de Hoon et al. 2005; Chen et al. 2013; Fleming et al. 2020; Georgakopoulos-Soares, Chan, et al. 2022; Miura, Ogake, and Ohyama 2018; von Hippel 1998). Therefore, we investigated whether the distribution of IRs is heterogeneous in functional genomic sites and whether they are preferentially positioned relative to TSSs and TESs in the genomes of organisms belonging to different taxonomic clades.

First, we examined the distribution of IRs across each genome’s functional genomic sub-compartments, including genome-wide, genic, exonic, 5’ UTR, 3’ UTR, and coding sequence (CDS) regions. We observed that, across the three domains of life and Viruses, IRs are predominantly concentrated in intergenic regions (**Figure 6A**). Notably, Bacteria exhibit the highest genome-wide IR density of 3,243.63 IR bps per Mb, followed by Viruses and Eukaryota (**Figure 6A**). Upon partitioning the various IRs by their spacer length, we noted significant differences in IR densities across the genomic compartments (**Figure 6B**). In Eukaryota, the exonic and CDS regions displayed low IR density for all spacer lengths, whereas a general propensity for perfect palindromes with zero bp spacer sequences is observed in genic regions. This leads us to conclude that IRs are favorably positioned within intronic regions and 3’ and 5’ UTR regions. This phenomenon is consistent across Plantae, Animalia, and Protista (**Supplementary Figure 6**-**7**). In Bacteria, the vast majority of IRs are positioned within intergenic regions with 4 bp being the most prevalent spacer length. In viral genomes, intergenic IR density is the highest; nonetheless, within non-coding regions, the predominant spacer length is 8 bp (**Figure 6B**; **Supplementary Figure 6**). When comparing individual phyla, we consistently observed that intergenic regions display the highest density of IRs across phyla, whereas there are larger differences in density at genic, exonic, and CDS regions between phyla (**Figure 6B**; **Supplementary Figure 6**-**7**). A subsequent analysis revealed that, when partitioning the various genes into protein-coding and non-coding groups, IRs are more prevalent in non-coding genes for Eukaryota and Archaea (Mann-Whitney U test, p-value<0.0001). Interestingly, for Bacteria, 4 bp was the most prevalent spacer length for IRs within non-coding genes (**Supplementary Figure 6**). These findings indicate that intergenic regions are most enriched for IRs across taxonomic groups, which could be due to the RNA polymerase stalling potential of hairpin and cruciform structures in genic regions (Hein et al. 2014).

**Figure 6:**
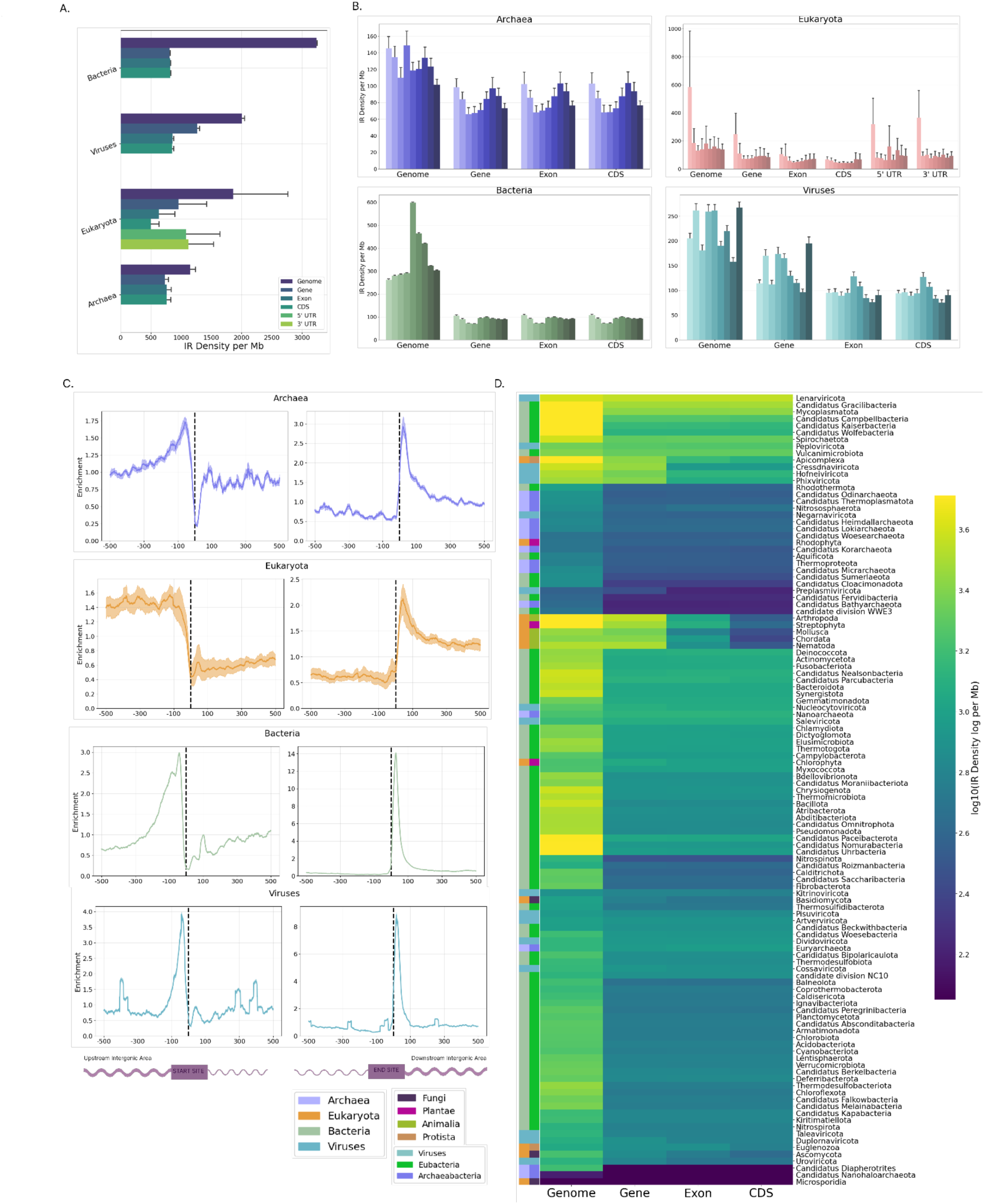
Examination of the topography of IRs across organismal genomes belonging to different taxonomies in the tree of life. **A.** Density of IRs across genomic subcompartments for the three domains of life and Viruses. **B.** Density of IRs across genomic subcompartments for the three domains of life and Viruses for the different spacer lengths. **C.** Density of IRs relative to the TSS and TES in the three domains of life and Viruses. **D.** Heatmap of the IR density across phyla at the CDS, exonic, genic, 5’ UTR, 3’ UTR (eukaryotic species), and genome-wide regions. Color bars on the left indicate the three domains of life and Viruses, as well as the six kingdoms to which the different phyla belong.

Next, we investigated whether IRs are differentially distributed relative to TSSs and TESs. We observed that across the three domains of life and Viruses, there is consistent enrichment of IRs in the 100-200 bp regions upstream of the TSSs, with peaks located at approximately -50 bp in Archaea, Bacteria, and Viruses, and at -150 bp in Eukaryota. IR densities drop dramatically at the TSSs (**Supplementary Figure 8**-**10**). IR motifs are also enriched in the regions downstream of the TESs in all domains and Viruses. IR densities exhibited very narrow and high peaks in Archaea, Bacteria, and Viruses within the tight regions of +50 bp to +80 bp downstream of the TESs (**Figure 6C**, **Supplementary Figure 10**-**12**), suggesting that they play roles in transcription initiation and termination across organismal genomes. Of particular note, the distribution of IRs after the TESs is 13-fold more enriched in Bacteria (**Figure 6C**) than in other domains and Viruses, which can be attributed to their role in rho-independent transcription termination (von Hippel 1998; d’Aubenton Carafa, Brody, and Thermes 1990). We found that the enrichment of IRs after the TESs in Bacteria is present across all phyla, albeit with varying effect sizes (**Figure 6C-D**, **Supplementary Figure 7,10**), indicating a general, ubiquitous bacterial mechanism of gene regulation. It is also noteworthy that the same distribution is present in Archaea, although with lower enrichment levels (**Figure 6C-D**, **Supplementary Figure 11**) and significant differences between archaeal phyla.

We segmented the IR counts into nine distinct groups characterized by the spacer length of the IR and studied the distribution of IR occurrences across the 1kb window around the TSS and TES for each spacer length. Furthermore, we partitioned the analysis to protein-coding and non-coding genes across the three domains of life, including viruses. In prokaryotic species, the protein-coding distribution of IR is enriched upstream of the TSS and downstream of the TES (**Supplementary Figure 13**). Notably, in prokaryotes, the non-coding distribution of IR is enriched downstream of the TSS; hence, the putative hairpin loop lies within the genic region and not upstream of the TSS. This discrepancy between non- coding and protein-coding IR positioning could be attributed to the different translational machinery deployed by the cell between the transcription of protein-coding versus non-coding RNAs, as well as the secondary structure of non-coding RNAs containing multiple hairpin loops. This phenomenon is not observed in eukaryotic organismal genomes, while in viruses the enrichment hotspots are dispersed in multiple regions, probably related to the different types of non-coding RNAs (**Supplementary Figure 13**). Additionally, in the TES, there is a higher degree of homogeneity between protein-coding and non-coding RNAs IR distribution across the four domains. In particular, in bacterial genomes, we found that the majority of IRs consisted of 4 bp spacers. Downstream of TES, we find that the IR hotspot consists primarily of IRs with 4 or 5 bp spacer length (**Supplementary Figure 13**). Experimentally this has been validated as the most stable forming hairpin loops (Varani 1995) and the stability of the hairpin loop could contribute to rho independent termination (Kingsford, Ayanbule, and Salzberg 2007; von Hippel 1998). These findings indicate that IRs are positioned relative to key regulatory elements and play roles in transcription initiation and termination across diverse organisms.

## Discussion

Here, we examined 118,065 complete organismal genomes to characterize the distribution, biophysical characteristics, and topography of perfect IRs. We observed that the frequency of perfect IRs varies substantially between different organismal genomes and taxonomic groups. Bacteria harbor the highest density of IRs, whereas Archaea have the lowest. However, large discrepancies are observed both between kingdoms and among different phyla within the same domain of life or Viruses. Such variation may suggest differences in environmental pressures, repair mechanisms associated with IRs, and/or the ability of certain organisms to process these structure-forming sequences in an error-free manner to mitigate the genomic instability associated with these sequences. For instance, replication fork stalling can occur at IRs (Voineagu et al. 2008; Lindsey and Leach 1989). In both prokaryotic and eukaryotic cells, IRs can cause genomic instability, including double-strand breaks, translocations, and deletions (Kirill S. Lobachev, Gordenin, and Resnick 2002; Narayanan et al. 2006; Hiroki Kurahashi et al. 2006; Lu et al. 2015). We also observed a subset of IRs with arms spanning tens of thousands of bps, which could have resulted from genomic rearrangement events such as inversions (Fried, Feo, and Heard 1991). Previous work has indicated that the presence of IRs is linked to a higher frequency of DNA double-strand breaks and inversions in human cells (Al-Zain et al. 2023); however, it remains to be studied whether similar mechanisms drive the formation of large IRs and whether hairpin structure formation at these large, perfect IRs contributes to genomic instability.

IRs were previously found to be substantially enriched in regions near start codons, stop codons, the ends of genes, 5’-untranslated regions, and promoter regions in *E. coli* (Miura, Ogake, and Ohyama 2018), consistent with our findings. These results suggest a role for IRs in gene expression regulation, potentially providing a selective evolutionary advantage that maintains IRs in these regions despite their intrinsic mutagenicity. In our study, although all domains and Viruses exhibited IR enrichments upstream of the TSSs and immediately downstream of the TESs, Eukaryota showed more variation, with wider enrichment peaks compared to other domains. This reflects the fact that eukaryotic cells generally have larger genes, longer intergenic regions, and more complex transcriptional control elements in their genomes.

The phyla studied exhibit diverse characteristics and show considerable variation in their IR genomic frequency, biophysical properties, and nucleotide composition. For instance, in prokaryotes, *Pseudomonadota* and *Bacillota* have lower genomic IR densities compared to *Mycoplasmatota*. Notably, *Aquificota*, autotrophs found in extreme environments such as hot springs and sulfur pools, display a significantly lower IR density compared to other bacterial phyla, similar to those of archaeal phyla. This could be due to specific environmental pressures affecting the likelihood of IR formation, as previously demonstrated (Tan and Chen 2008), and potentially negatively impacting the organism’s ability to resolve them. The preferential distribution of IRs relative to TSSs and TESs was particularly elevated in Bacteria and Archaea. This can be explained by the role of IRs in rho-independent transcription termination (Kingsford, Ayanbule, and Salzberg 2007; von Hippel 1998). The presence of dyad symmetry in broader terminator regions halts the transcription process by enabling the dissociation of the ternary elongation complex, causing the mRNA polymerase to detach. This phenomenon is widespread across bacterial species, where IRs play a vital role in the intrinsic termination process.

Furthermore, the enrichment of IRs relative to regulatory elements, including TSSs and TESs, indicates that IRs are not merely statistical artifacts but are essential for key regulatory and functional mechanisms within the cell, reinforcing previous research (Georgakopoulos-Soares, Victorino, et al. 2022; Siegel et al. 2022; Brázda et al. 2020; de Hoon et al. 2005; Chen et al. 2013; Fleming et al. 2020; Georgakopoulos-Soares, Chan, et al. 2022; Miura, Ogake, and Ohyama 2018; Kingsford, Ayanbule, and Salzberg 2007). Additionally, the differences in IR densities between taxa may reflect functional roles specific to individual taxonomic groups, as previously reported (Kingsford, Ayanbule, and Salzberg 2007; Bowater, Bohálová, and Brázda 2022; Poggi and Richard 2021).

We conclude that IRs are highly plastic genomic elements, displaying significant variation in frequency across different taxonomic subgroups and within organisms across functional elements.

## Methods

### Data retrieval and parsing

Complete organismal genomes were downloaded from the GenBank and RefSeq databases (O’Leary et al. 2016; Benson et al. 2013). Duplicate assembly accessions were filtered out. Gene annotation files in the form of GFF files were downloaded for each genome from the same source. Coordinates for genes, exons, and CDS regions were derived using BEDTools (Quinlan and Hall 2010) and custom awk, bash, and Python scripts. These coordinates were then examined for IR densities across organismal genomes. TSSs and TESs were derived from the corresponding gene coordinates.

### Identification of IRs in organismal genomes

IRs with arm lengths equal to or longer than ten bps, spacer lengths less than nine bps, and without mismatches in the arms were used throughout the study. For IR detection, a modified version of the non-B gfa package was developed and wrapped in a Python program (Cer et al. 2013). The subprocess call to the C script utilized the parameters *minIRrep=8* and all necessary skip flags to extract only the IR sequences present in each organismal genome. A custom script was used to manually transform the data into a processable tabular format and extract the corresponding arm and spacer sequences along with their corresponding lengths based on the coordinates of the IR sequence. The detected IRs were then systematically validated.

### Examination of IR frequency across taxonomic subdivisions

Organismal genomes were subdivided into different taxonomic levels, namely the three domains of life and Viruses, as well as kingdoms and phyla. The distribution and number of IRs were examined between taxonomic groups at the same taxonomic level by aggregating organisms within each taxonomic group. The density of IRs was defined as the number of IR bps divided by the total bps in a genome or genomic sub-compartment, multiplied by 1,000,000 to present the density in IR bps/Mb. For each domain, kingdom, and phylum, the average IR density (Mb) of the associated organismal genomes was calculated. Error bars were determined using the standard error.

### Examination of biophysical properties of inverted repeats

The number of IRs identified was calculated separately for each combination of spacer and arm length, and the proportions were compared between different taxonomic groups. The mononucleotide and dinucleotide compositions, as well as the GC and AT content of spacers and arms of IRs, were calculated and examined for different arm and spacer lengths. These comparisons were performed across different taxonomic levels.

Using the Jellyfish program (Marçais and Kingsford 2011), we counted the occurrences of mono- and di-nucleotides in IR spacers and arms, as well as across each organismal genome. We then performed the enrichment comparison between the IR frequency and the genome frequency of the different mono- and dinucleotides. Word cloud plots were constructed by including only IRs that appeared exclusively in one of the four domains, in order to provide insights specific to that domain.

The Venn Diagram in Figure 1d was generated by including only the unique IR arms present in each domain of life or across Viruses. For the construction of the upset plot, unique IRs were partitioned into 15 mutually disjoint sets, and the total number of unique IRs was calculated for each set. Furthermore, each unique IR was recorded with the specific set of spacer lengths it was associated with, and this was divided by the total number of possible spacer lengths - nine in our case - allowing the calculation of the spacer length diversity index. This index represents the percentage of possible spacer length configurations with which the sequence was found across the various organismal genomes.

### Inverted repeat topography across functional genomic sub-compartments

For each organismal genome, the density of IRs in intergenic, genic, exonic, 5’ UTR, 3’ UTR, and CDS regions was calculated using publicly available GFF coordinate files. A workflow program was developed to process and handle subtle inconsistencies in the downloaded GFF files. To estimate the total IR coverage, missing transcripts and exonic regions of protein-coding genes from prokaryotic assemblies were filled in for each GFF file. Since 3’ and 5’ UTR regions are not present by default, a custom script was used to derive these regions. Subsequently, for each annotated subcompartment, overlapping compartments of the same type were merged and expanded into non-overlapping, mutually disjoint sets of coordinates. These coordinates were then used to estimate the total coverage per Mb in each organismal genome by calculating the total number of bps of IRs divided by the total compartment length of the previously merged genomic regions. Finally, all aggregated statistics for each organismal genome were concatenated into a large, snappy-compressed parquet file, which was then processed to generate the depicted figures.

The density of IRs was calculated at each position within a window relative to the TSS and TES. Enrichment was estimated as the number of IR bps at a position divided by the mean number of bp occurrences across the window of the distribution, as previously described (Georgakopoulos-Soares, Parada, et al. 2022). Confidence intervals in the distribution plots were calculated as the standard error at each position.

The IR density in genic, exonic, and CDS regions (Mb) was calculated by dividing the raw total number of IR overlap bps by the total compartment length in bps and then multiplying the resulting density by 1,000,000 (Mb). The intergenic IR density (Mb) was calculated by subtracting the total number of IR bps found at a given organismal genome from those IR bps at genic regions. This result was then divided by the total intergenic length of the genome.

The enrichment density plots across the tree of life were generated by expanding the window size by 500 bps upstream and downstream from the TSS and TES. For each organismal genome, an interval of 1001 bps was constructed and summed symmetrically to determine the total number of IR bps at each coordinate, with 0 representing either the TSS or TES. Each genomic interval was divided by the window mean to account for differences in genome size, as previously described (Georgakopoulos-Soares, Parada, et al. 2022). Finally, for every domain, each genomic location was averaged across all organismal genomes to generate the enrichment density plots.

## Statistical analyses

Statistical analyses were performed in Python using the libraries Scipy (Virtanen et al. 2020), and NumPy (Harris et al. 2020).

## Code Availability

The code to perform these analyses can be found at: https://github.com/Georgakopoulos-Soares-lab/inverted_repeats

---

**Supplementary Figure 1:**
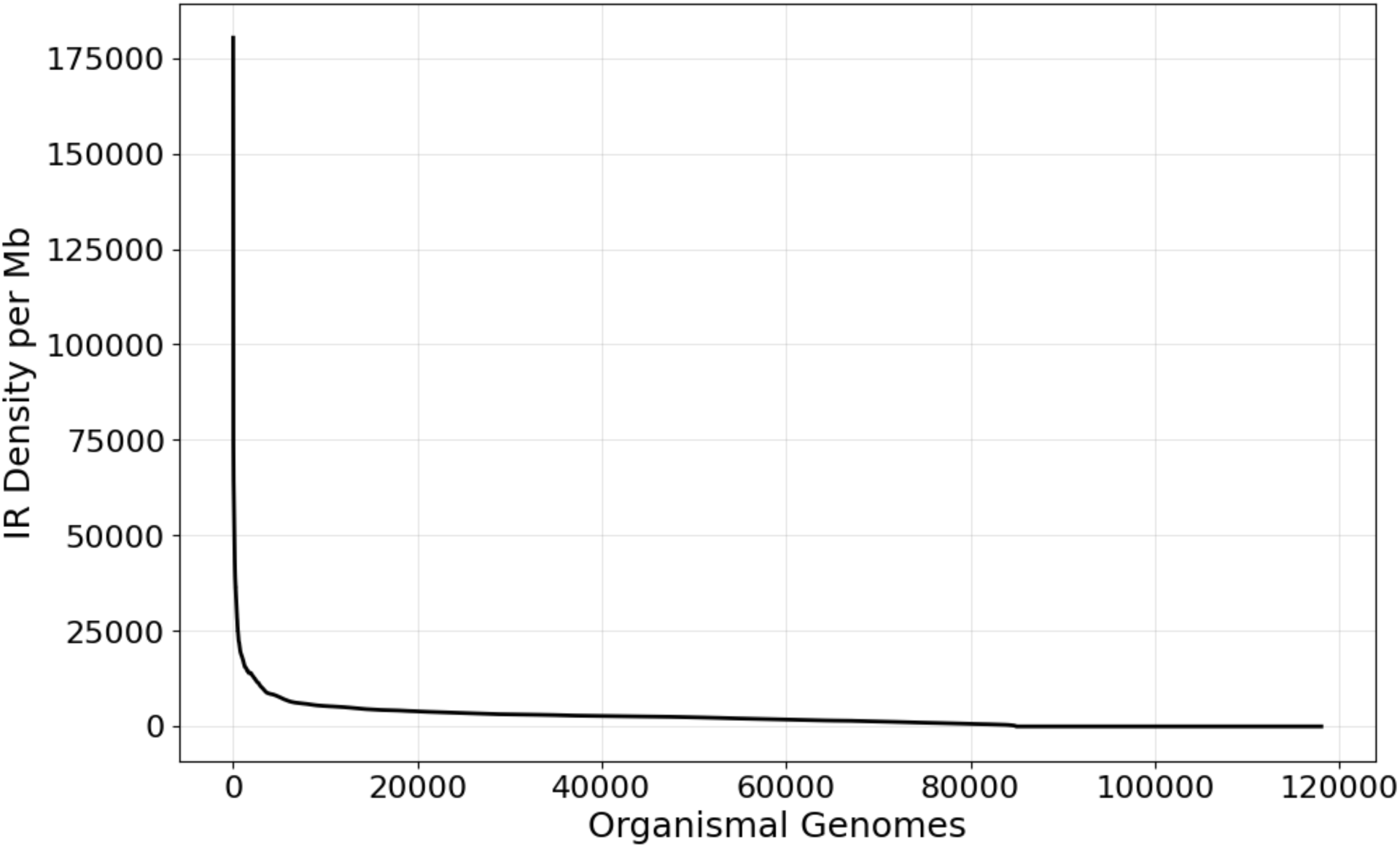
Distribution of the IR density per Mb across the organismal genomes studied.

**Supplementary Figure 2:**
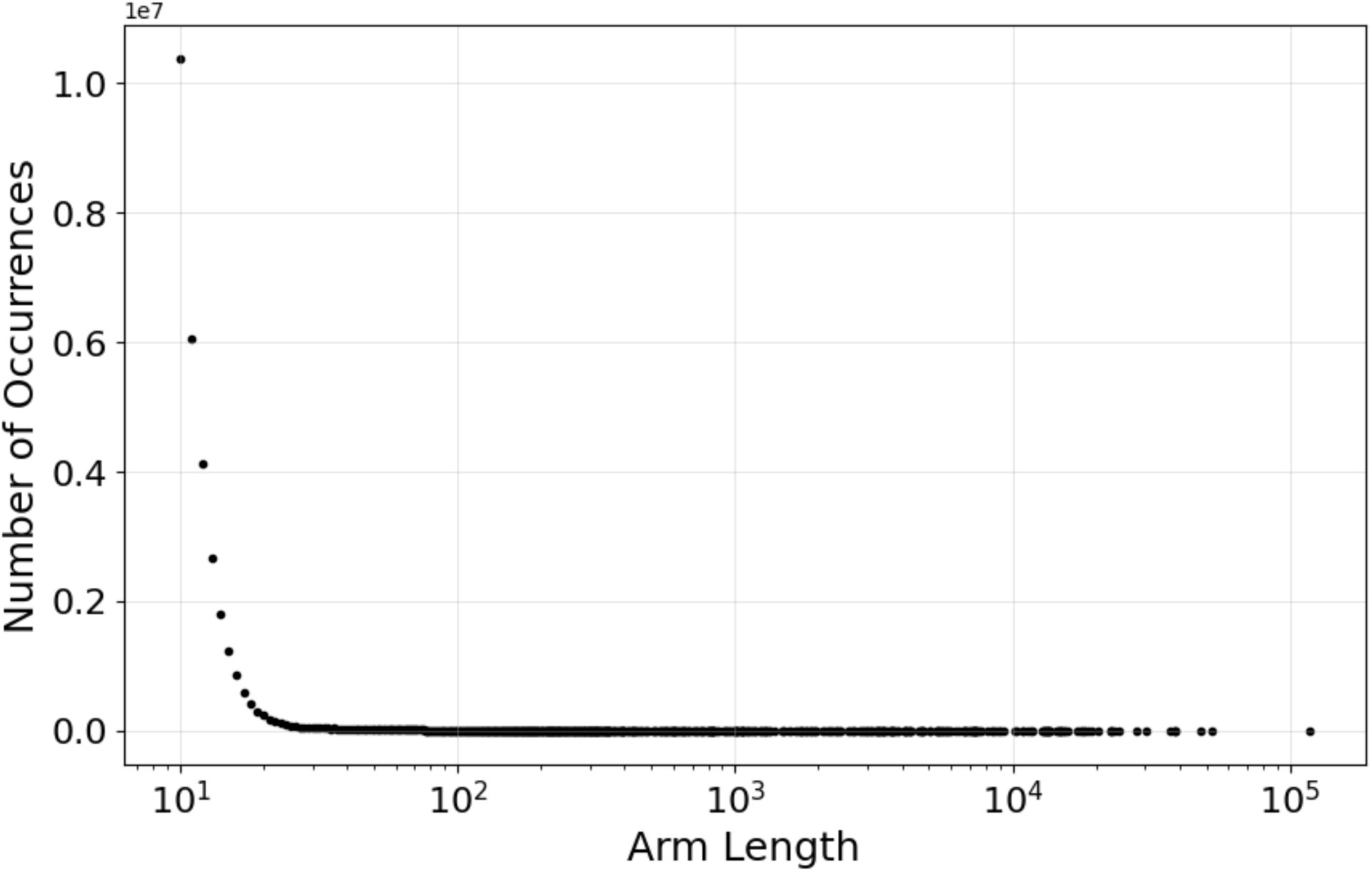
Number of IRs found for different arm lengths.

**Supplementary Figure 3:**
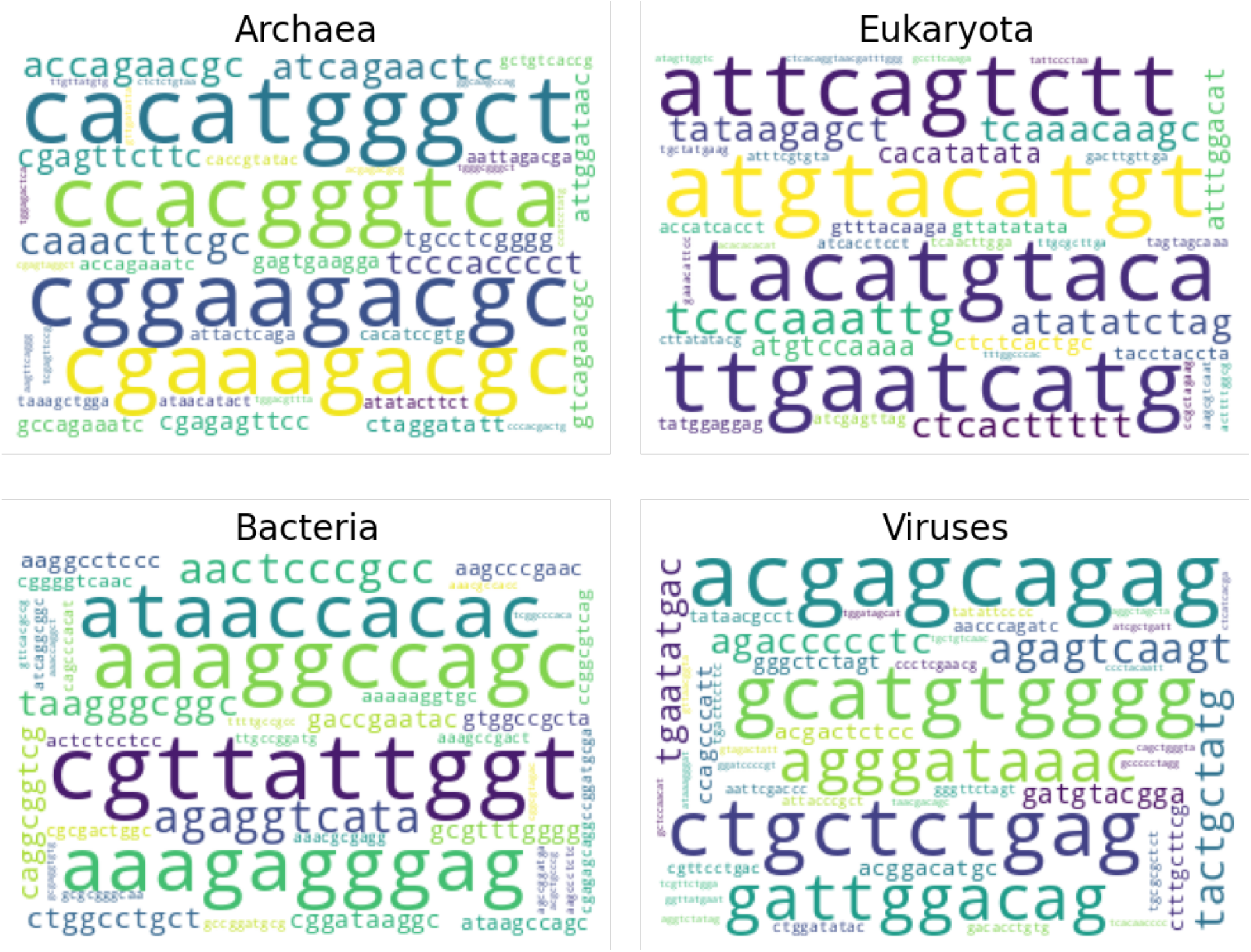
Unique IR arms in each of the three domains of life and Viruses. Results shown for arm length of ten bps.

**Supplementary Figure 4:**
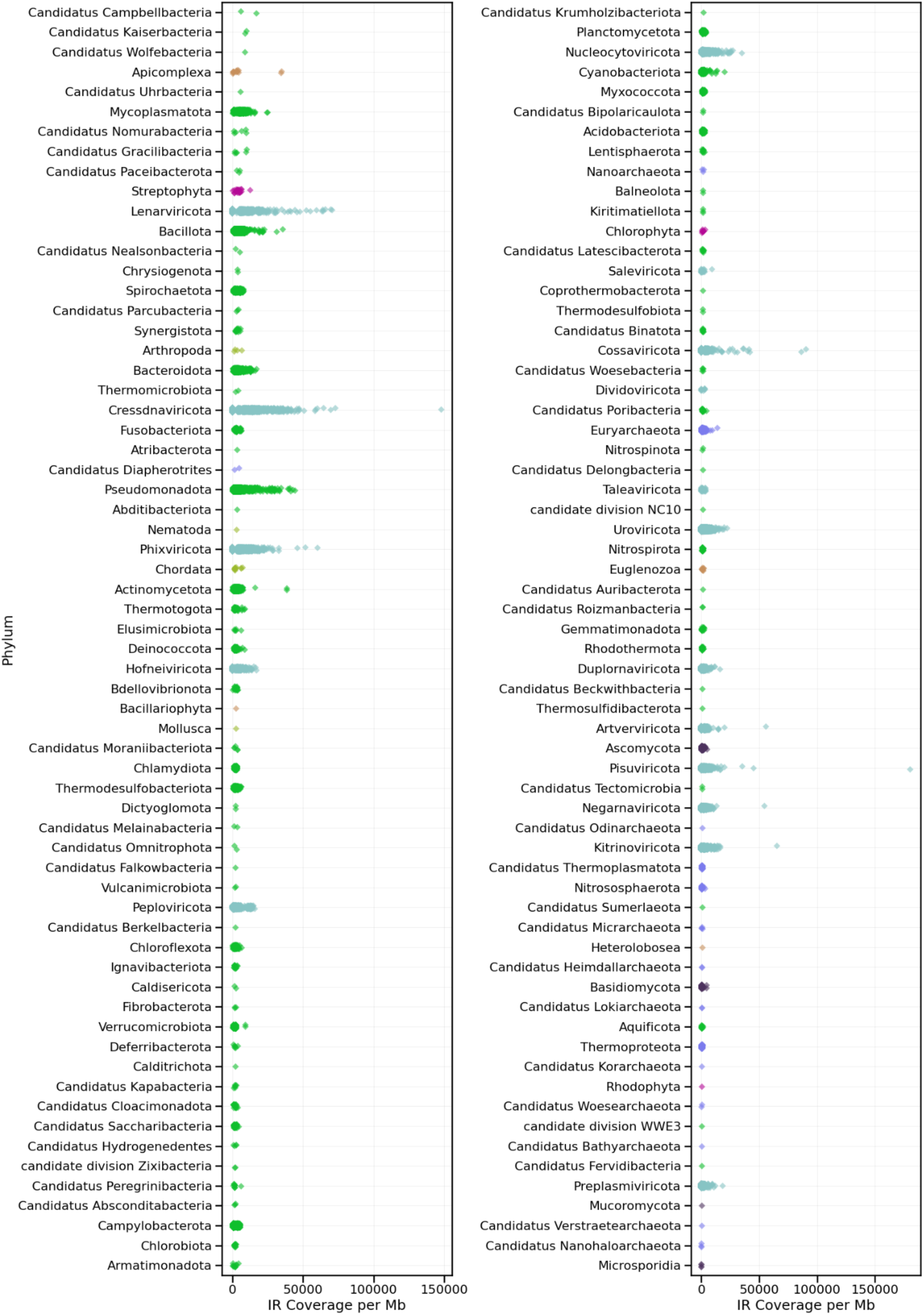
IR coverage across organismal genomes in the different phyla. Every dot represents a species, and colors represent the different kingdoms.

**Supplementary Figure 5:**
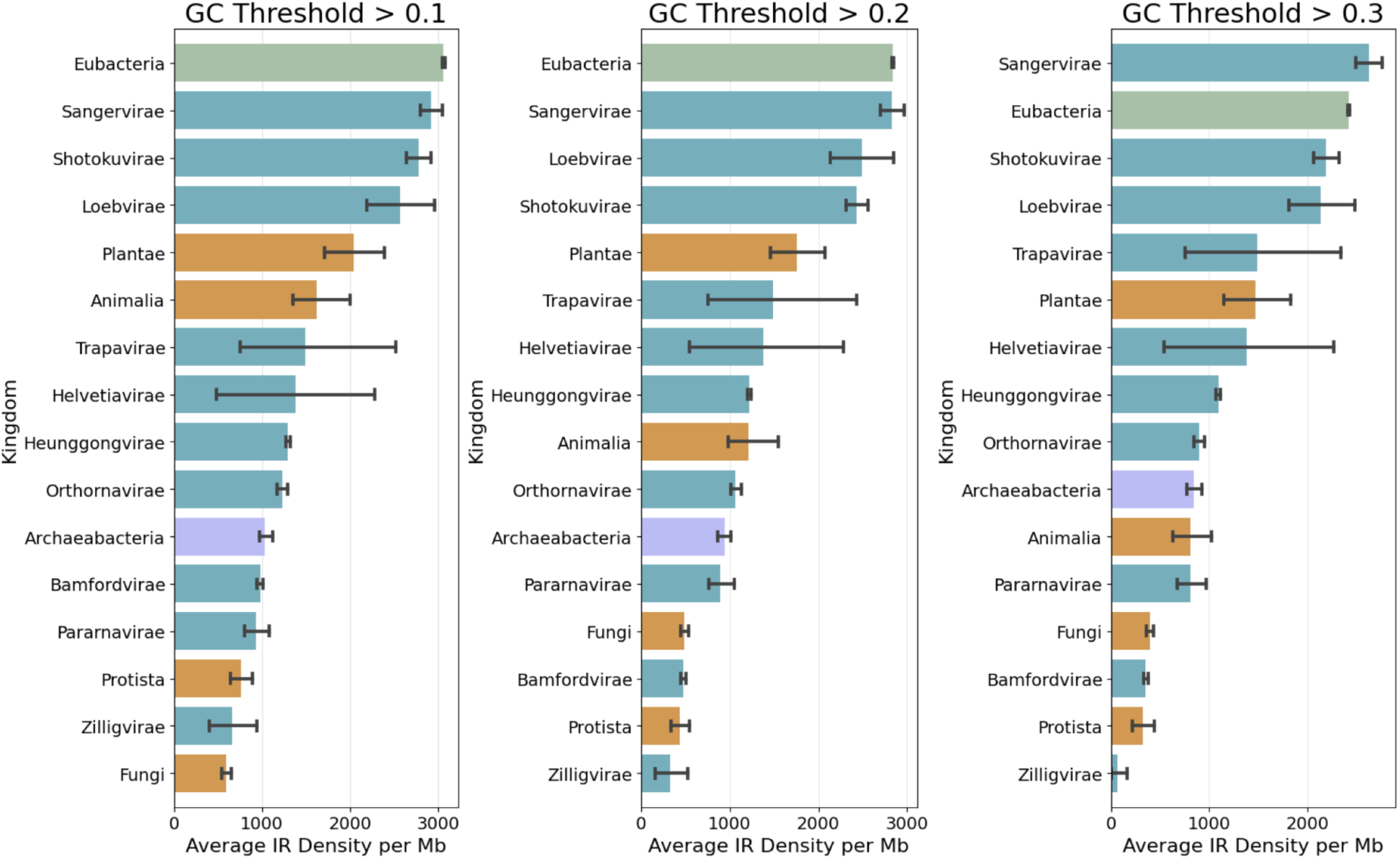
IR coverage across organismal genomes in the different kingdoms for different thresholds of the GC content of the IR. Results shown for GC% higher than: a. 10%, b. 20%, c. 30%.

**Supplementary Figure 6:**
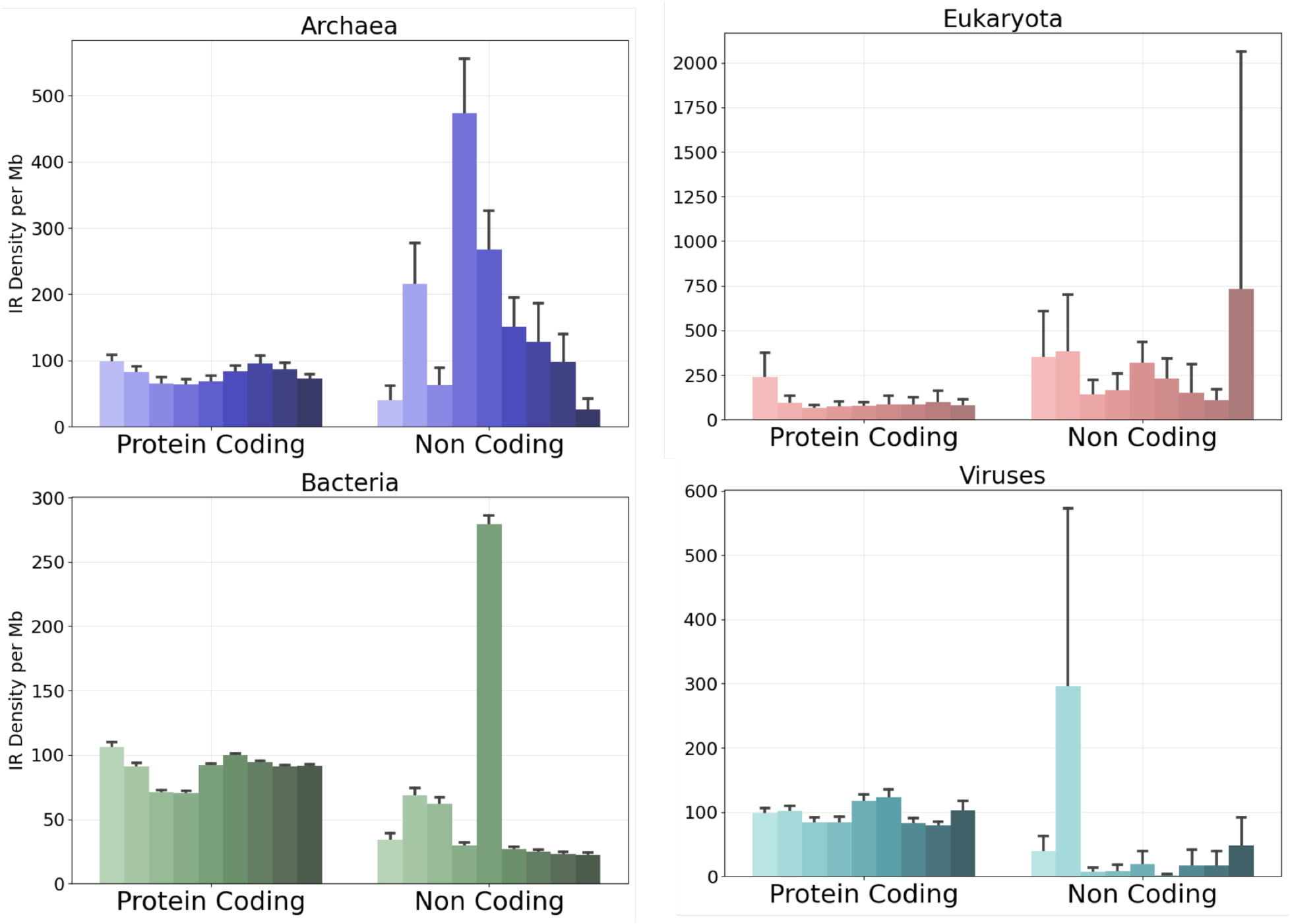
Comparison of IR density separated by gene type into coding- and non-coding and by spacer length across the three domains of life and Viruses.

**Supplementary Figure 7:**
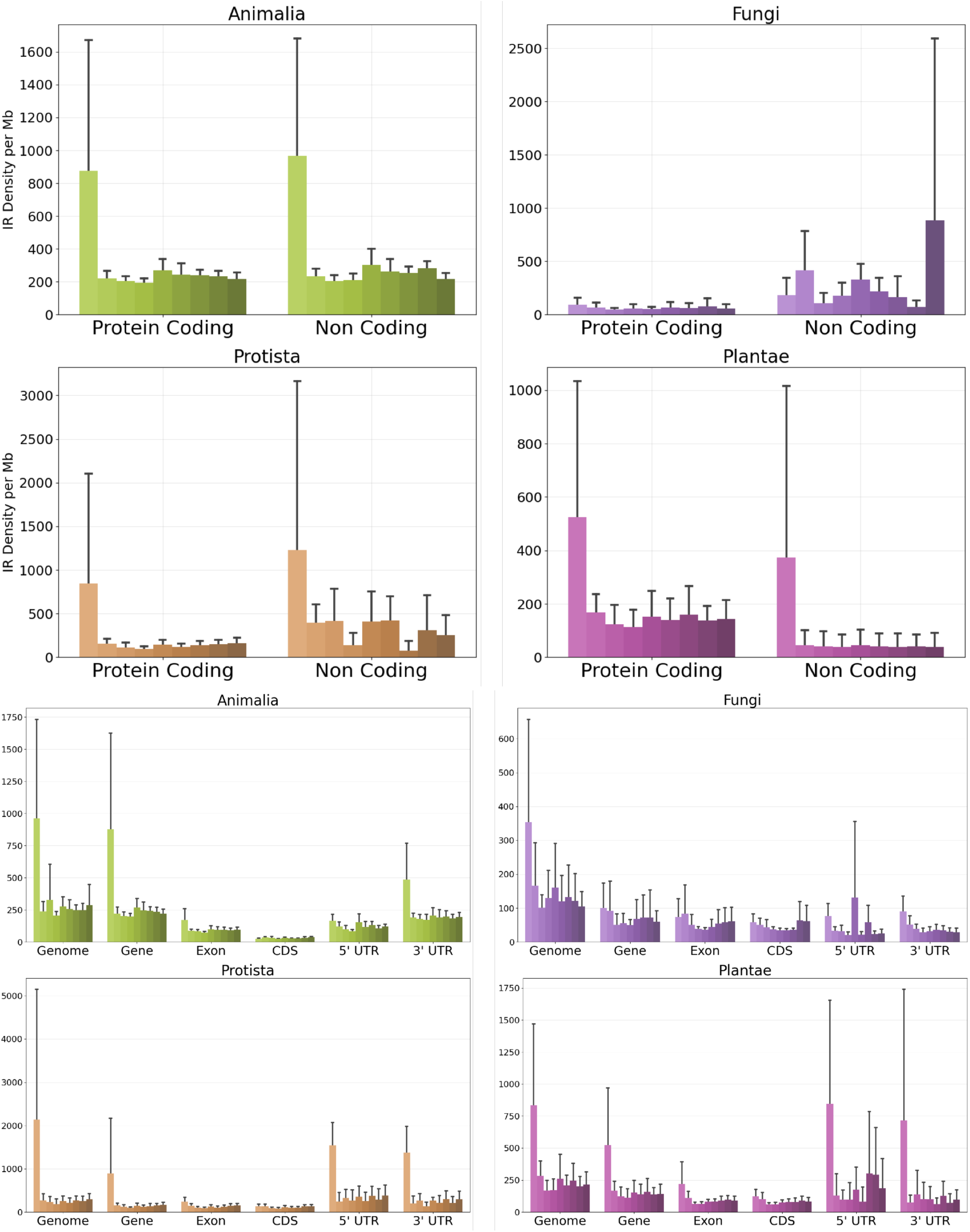
Comparison of IR density separated by gene type into coding- and non-coding and by spacer length across eukaryotic kingdoms.

**Supplementary Figure 8:**
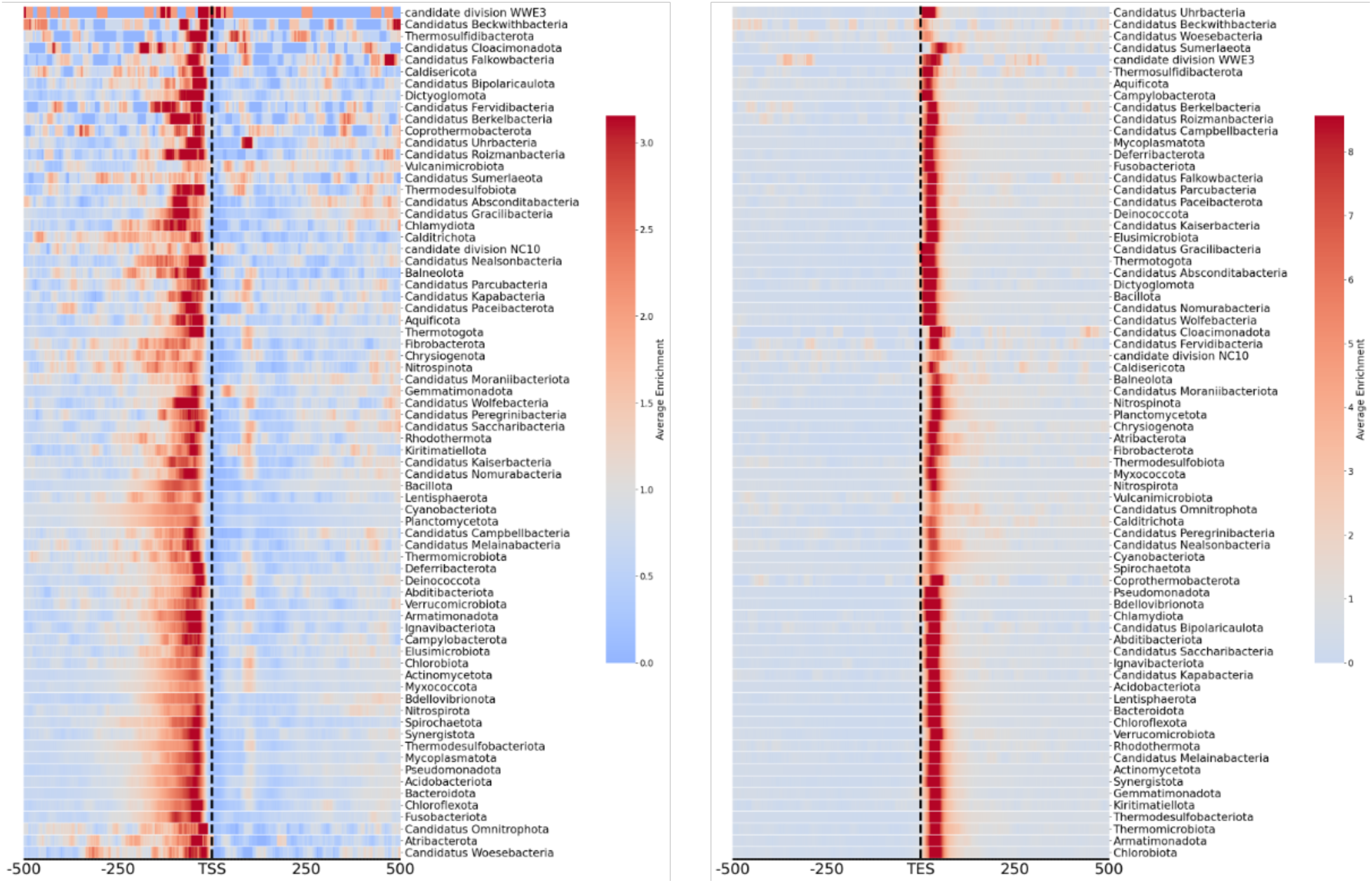
Enrichment of IRs relative to the TSS and TES in bacterial phyla.

**Supplementary Figure 9:**
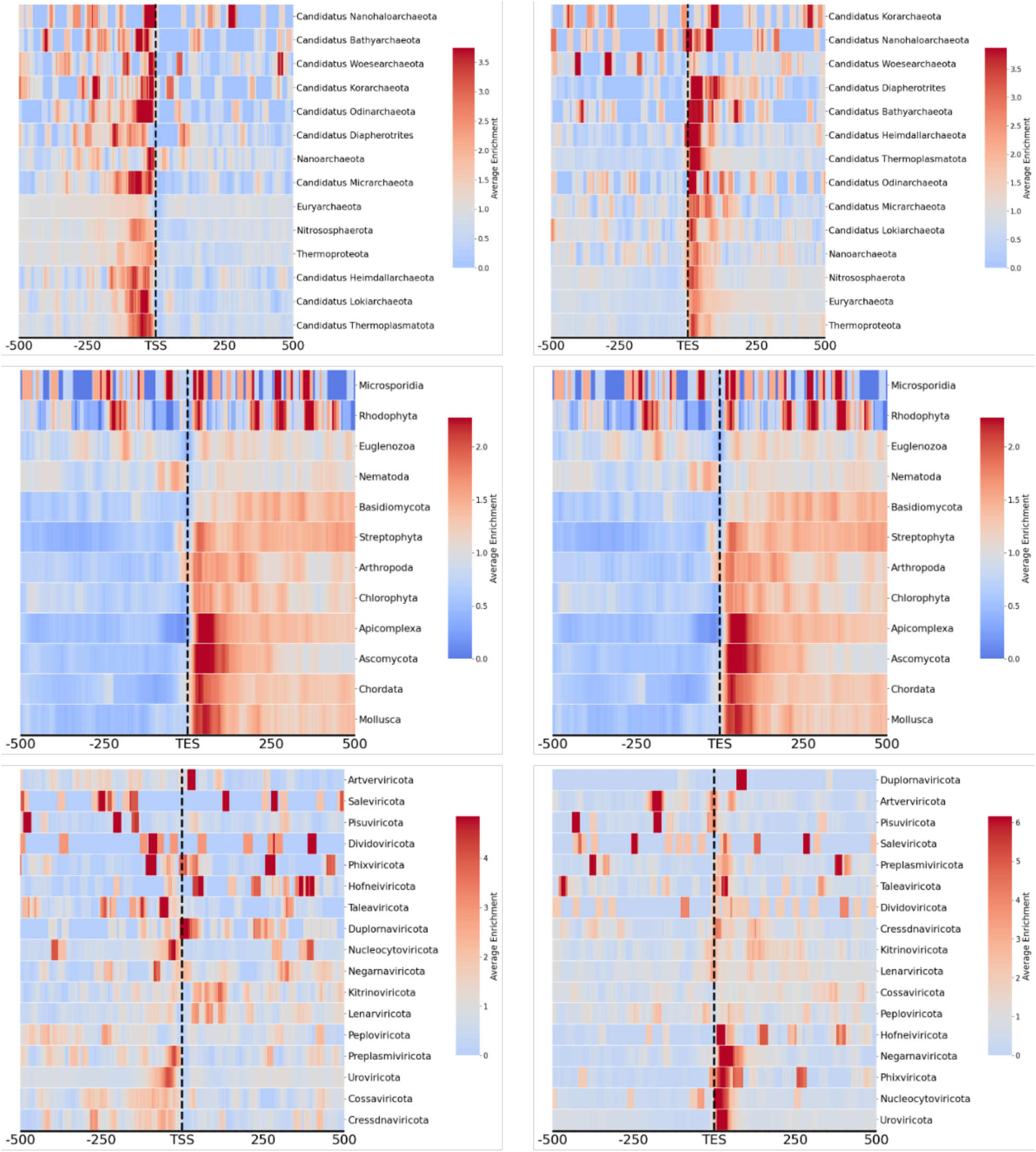
Enrichment of IRs relative to the TSS and TES in archaeal, eukaryotic, and viral phyla.

**Supplementary Figure 10:**
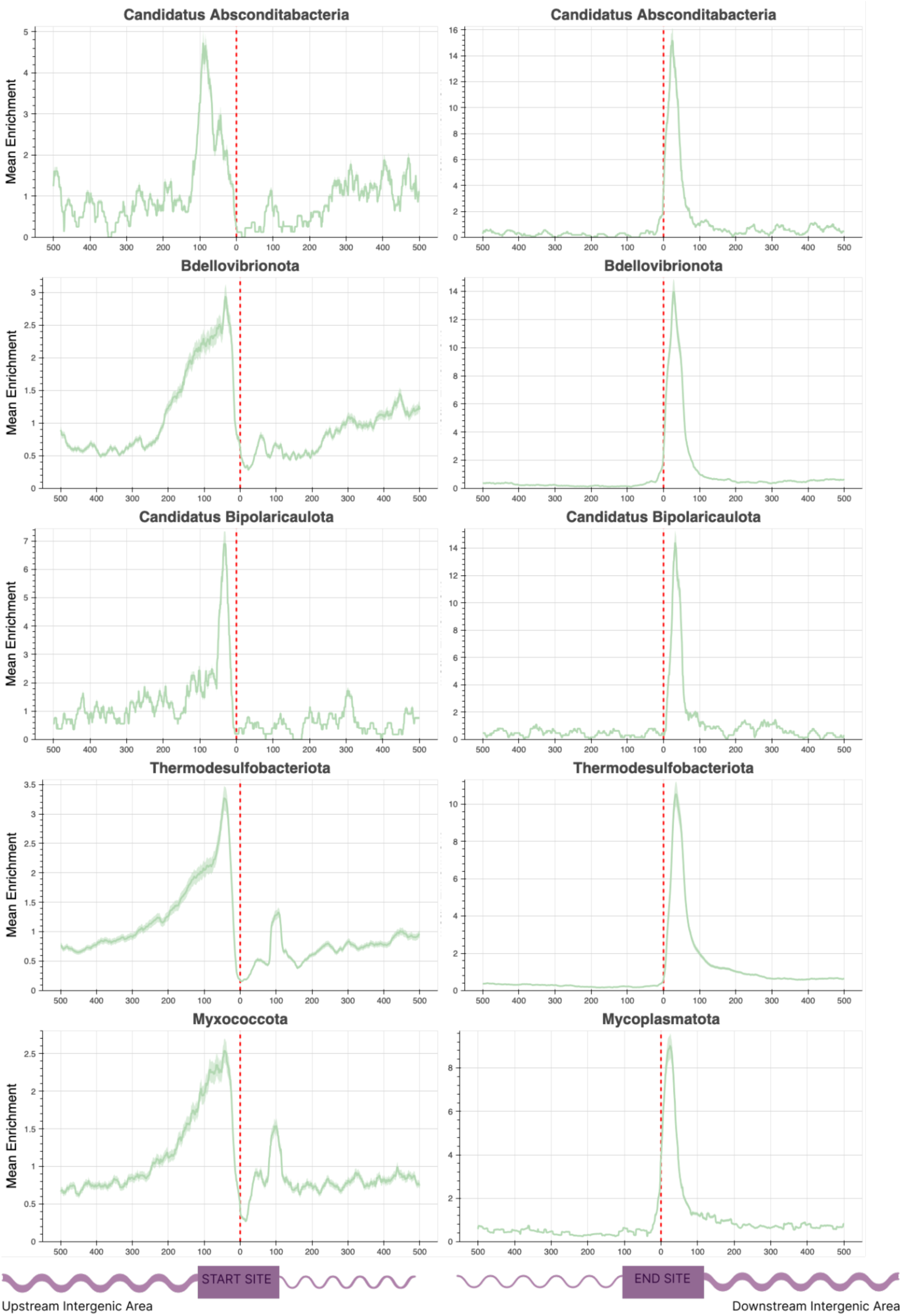
Enrichment of IRs relative to the TSS and TES in individual bacterial phyla.

**Supplementary Figure 11:**
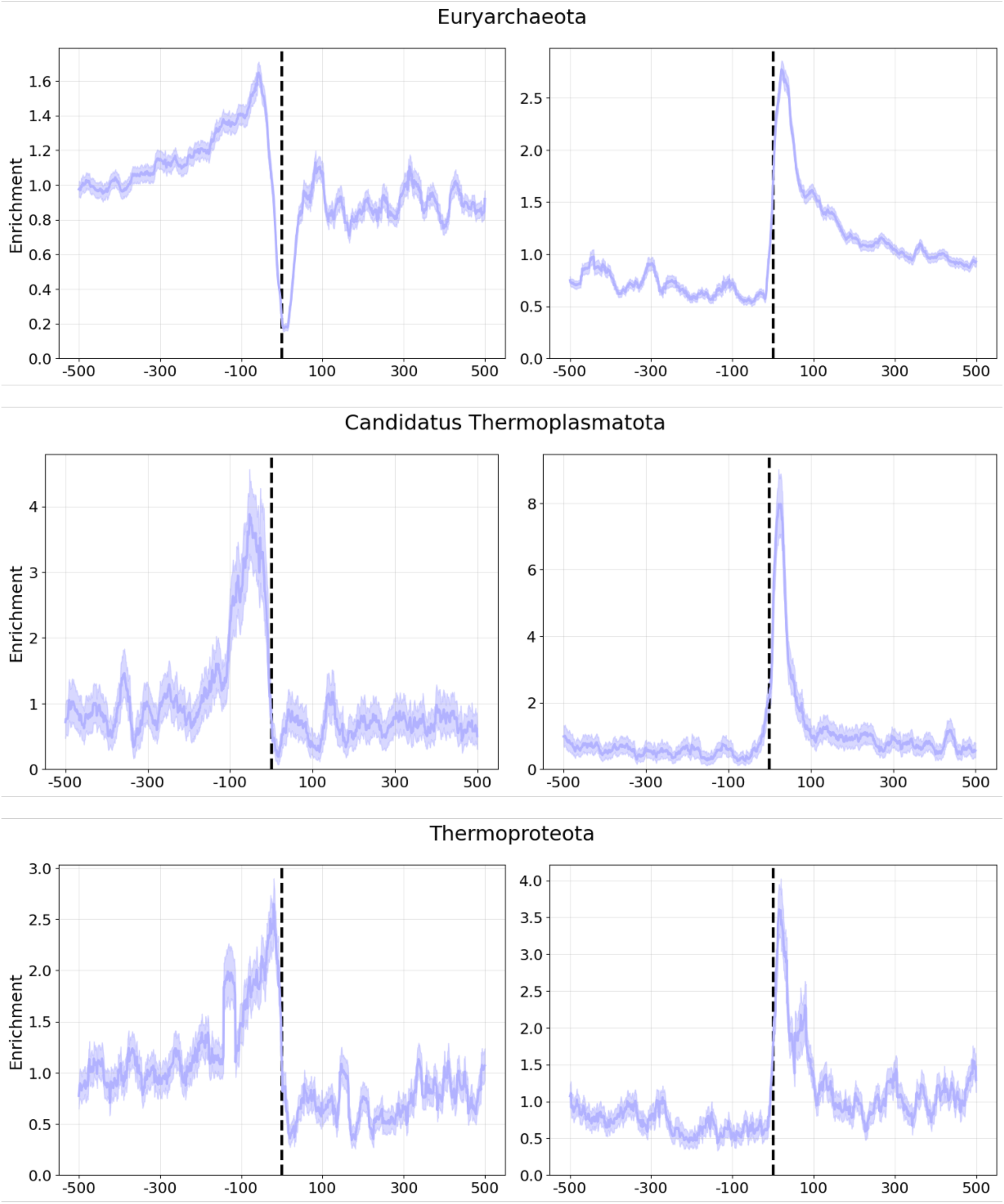
Enrichment of IRs relative to the TSS and TES in individual archaeal phyla.

**Supplementary Figure 12:**
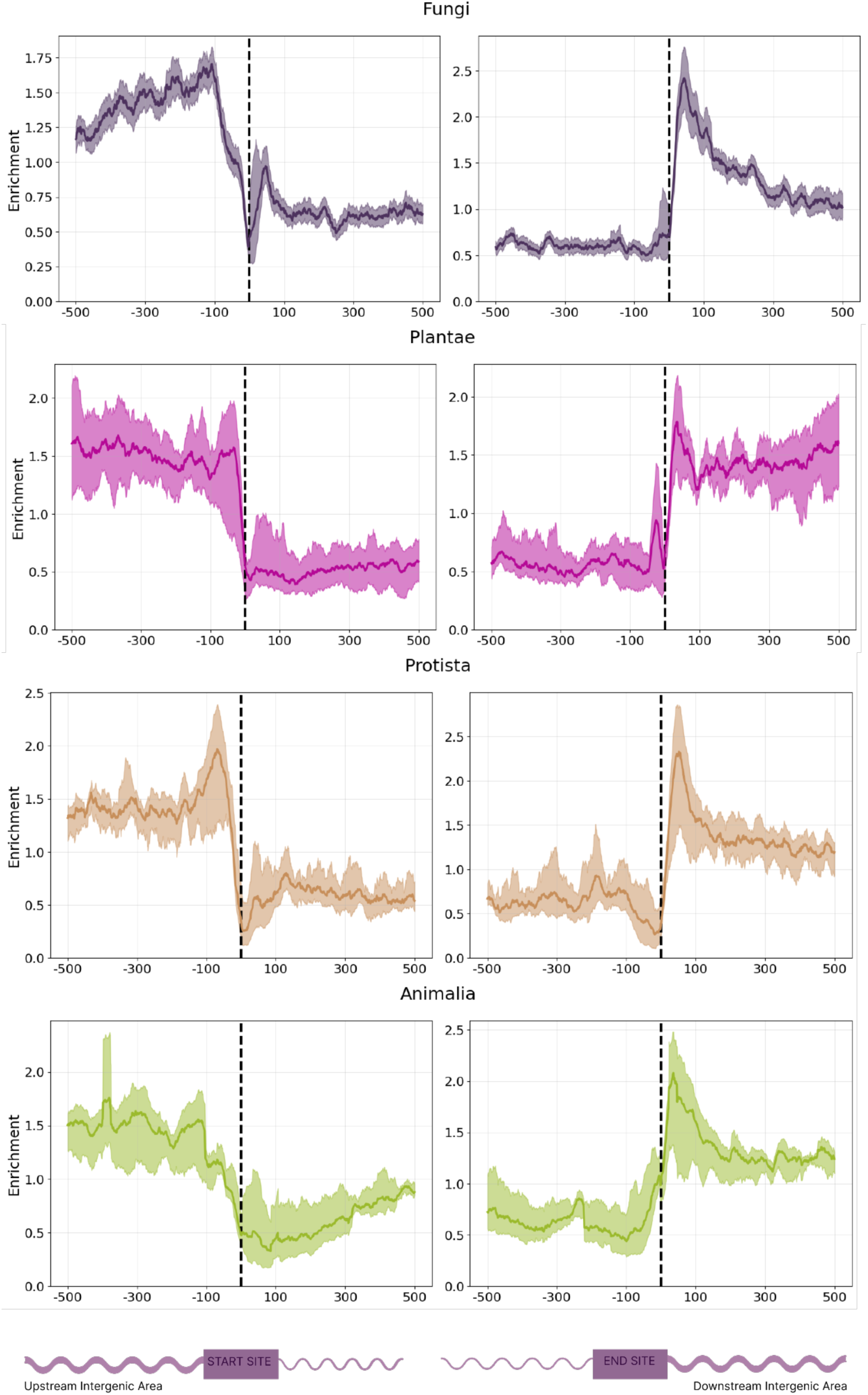
Enrichment of IRs relative to the TSS and TES in individual eukaryotic kingdoms.

**Supplementary Figure 13:**
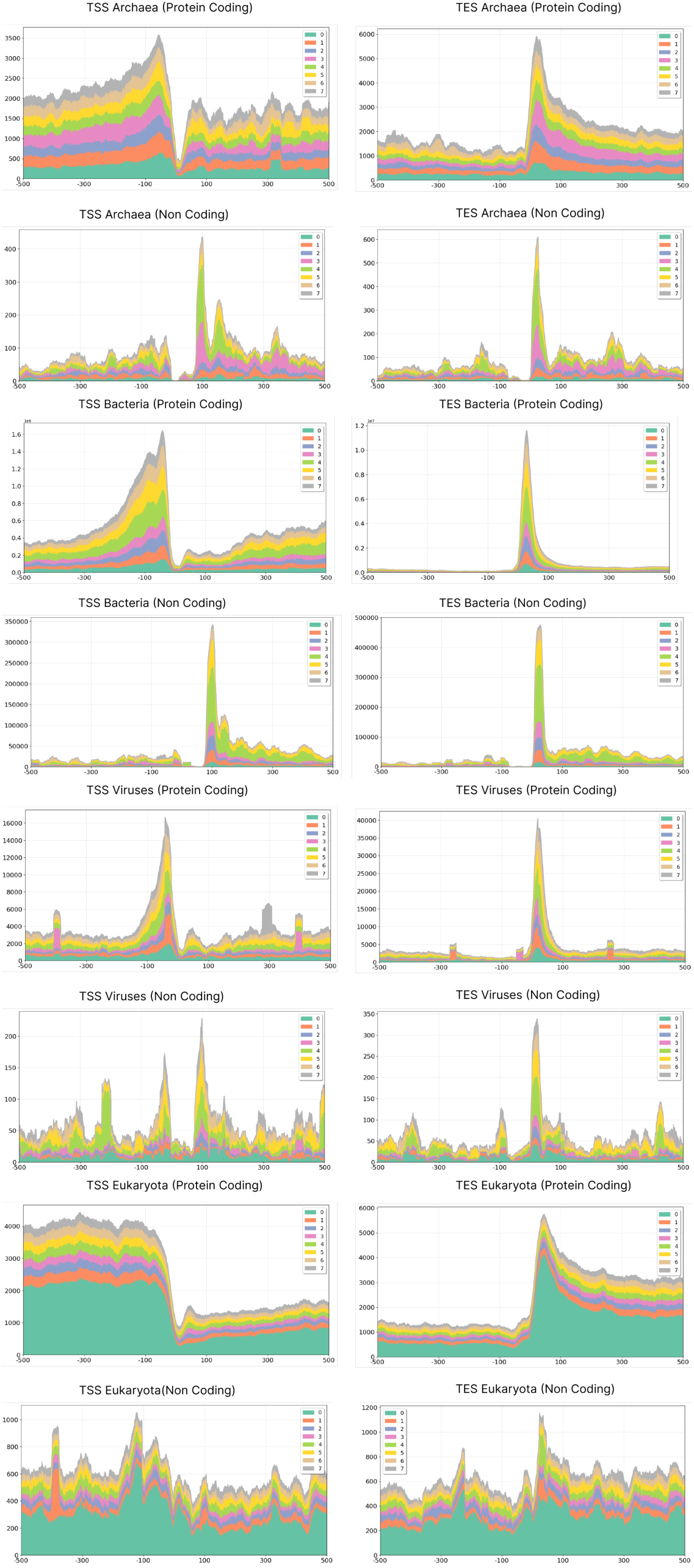
Distribution of IRs separated by spacer length, in coding- and non-coding genes.

**Supplementary Table 1:** Largest perfect IRs detected in *Homo Sapiens*.

| Chromosome | Start (bp) | End (bp) | Arm Length (bp) | Spacer Length (bp) | GC Content (%) |
| --- | --- | --- | --- | --- | --- |
| chr8 | 125,615,789 | 125,616,077 | 140 | 8 | 2.08 |
| chr8 | 87,939,831 | 87,940,074 | 118 | 7 | 1.23 |
| chr9 | 114,631,600 | 114,631,798 | 95 | 8 | 25.75 |
| chr11 | 42,828,571 | 42,828,757 | 93 | 0 | 18.28 |
| chrX | 138,732,619 | 138,732,799 | 88 | 4 | 37.22 |
| chr3 | 60,903,975 | 60,904,152 | 86 | 5 | 11.86 |
| chr6 | 67,938,601 | 67,938,769 | 84 | 0 | 15.48 |
| chr11 | 5,253,730 | 5,253,890 | 80 | 0 | 37.50 |
| chr2 | 85,010,163 | 85,010,327 | 80 | 4 | 6.71 |
| chr13 | 83,726,139 | 83,726,299 | 80 | 0 | 3.75 |

## Notes

### Competing Interest Statement

The authors have declared no competing interest.

